# Transiently elevated dopamine triggers plasticity that preserves motor function, striatal neuron morphology, and thalamostriatal input in a mouse model of Parkinson’s disease

**DOI:** 10.1101/2024.05.28.596192

**Authors:** Joe C. Brague, Ghanshyam P. Sinha, David A. Henry, Daniel J. Headrick, Zane Hamdan, Bryan M. Hooks, Rebecca P. Seal

## Abstract

Loss of nigrostriatal dopamine (DA) neurons in Parkinson’s disease (PD) produces motor symptoms by disrupting the normal circuitry of striatal projection neurons. Here, we report that the Transient Elevation of Dopamine (TED) within a defined window of 6-OHDA-induced DA depletion preserves normal motor behavior for months. We also observed a preservation of mature spines specifically on direct pathway striatal projection neurons (dSPNs) and not indirect pathway striatal projection neurons (iSPNs). Preservation of motor behavior and dSPN spines required enhanced activation of striatal D1 receptors. Circuit mapping showed input from primary motor cortex was reduced, while input from parafascicular thalamus was unchanged in both TED mice and controls of both sexes, suggesting rewiring of excitatory inputs to dSPNs from other cortical or thalamic areas in TED mice. Lastly, thalamostriatal paired pulse facilitation, normally depressed following DA depletion, is preserved in TED mice. Although the causal basis of motor preservation remains unresolved, our findings reveal that a schedule of transiently increasing striatal DA surrounding the depletion of dopamine can engage a long-lasting plasticity that markedly delays the onset of motor impairment.

**SIGNIFICANCE STATEMENT:** In Parkinson’s disease, death of dopamine-producing neurons disrupts brain regions that coordinate movement, leading to stiffness and slowness. Available therapies relieve symptoms by supplementing the lost dopamine. Here we show that increasing dopamine neuron activity in two genetic mouse models for a brief period primes the circuit to adapt, delaying the onset of motor impairment (for months) despite a severe loss of dopamine. The mechanisms that underlie this adaptation could open new paths to saving off debilitating motor impairments in Parkinson’s patients.

## INTRODUCTION

The dorsal striatum is critical for voluntary movement. The major output neurons, GABAergic spiny projection neurons (SPNs), comprise >95% of striatal neurons and receive dense glutamatergic input from the cortex and thalamus as well as dopaminergic (DA) input from the substantia nigra pars compacta (SNpC). Direct pathway SPNs (dSPNs) express the D1 dopamine receptor and project to the substantia nigra pars reticulata (SNpR) and globus pallidus internal, whereas indirect pathway SPNs (iSPNs) express the D2 receptor and project to the globus pallidus external and subthalamic nucleus (Albin, Young et al. 1989, Gerfen, Engber et al. 1990, Smith, Bevan et al. 1998, Villalba and Smith 2010, Lanciego, Luquin et al. 2012, Villalba and Smith 2018, McGregor and Nelson 2019). The output of these pathways influences voluntary movement control, including initiation, speed, and cessation, through mechanisms that are not entirely understood; however the direct pathway plays a role in facilitating movement (Kravitz, Freeze et al. 2010, Cui, Jun et al. 2013, Panigrahi, Martin et al. 2015, Markowitz, Gillis et al. 2018). dSPNs and iSPNs receive two major sources of long-range excitatory input: corticostriatal projections arising from several cortical regions (Alexander, DeLong et al. 1986, Hintiryan, Foster et al. 2016), most prominently primary motor cortex (M1), and thalamostriatal input mainly from intralaminar nuclei (Smith, Raju et al. 2009).

In Parkinson’s disease (PD), the loss of DA input from SNpC to the dorsal striatum impairs voluntary movement through changes in the activity of SPNs, shifting striatal output to favor iSPNs (Stephens, Mueller et al. 2005, Smith, Raju et al. 2009). These changes manifest as altered excitatory connectivity to SPNs and altered intrinsic excitability. An increase in the intrinsic excitability of dSPNs is observed consistently across rodent models, while changes in iSPNs are more variable across models (Fieblinger, Graves et al. 2014, Maurice, Liberge et al. 2015). Consistent with altered excitatory input, mature spines and dendritic arbors of both SPN types are reduced in 6-OHDA rodent models of PD (Day, Wang et al. 2006, Fieblinger, Graves et al. 2014, Gagnon, Petryszyn et al. 2017, Villalba and Smith 2018), the MPTP primate model (Smith and Villalba 2008, Smith, Raju et al. 2009, Villalba and Smith 2010) and postmortem brains of PD patients (Stephens, Mueller et al. 2005, Zaja-Milatovic, Milatovic et al. 2005). Thus, altered excitatory input to the striatum is thought to contribute to movement impairment in PD and rebalancing input to SPNs can restore motor function. For example, in the median forebrain bundle (mfb) 6-OHDA mouse model, Parker et al showed that cortical input to both dSPNs and iSPNs was similarly reduced, whereas thalamic input from the parafasicular nucleus (PF) was increased to iSPNs and decreased to dSPNs (Parker, Lalive et al. 2016). Decreasing the strength of the thalamostriatal output from PF neurons after DA depletion restored normal motor function. Thus, strategies that alter SPN extrinsic inputs to rebalance striatal output, can be an effective strategy to alleviate PD motor deficits.

We identified transgenic mice that show normal motor behavior following dorsal striatal 6-OHDA-induced DA depletion (Divito, Steece-Collier et al. 2015). The preserved motor behavior lasts for up to 20 weeks. Under non-depleted conditions, vesicular glutamate transporter (VGLUT) 3 knockout mice show a circadian-dependent increase in striatal DA release, locomotion, and immature spines during the waking cycle (Divito, Steece-Collier et al. 2015). This study investigates whether the daily increase in DA release we observed in VGLUT3 KO mice underlies the preserved motor function. We report that preservation of motor behavior similarly occurs in an inducible mouse model in which endogenous striatal DA was elevated to mimic the increase in striatal dopamine of the VGLUT3 KO using an excitatory designer receptor activated by designer drugs (eDREADD) expressed in midbrain DA neurons. Induction of preserved motor behavior requires DA neurons are activated for at least 3 days prior to and 5 days during depletion. Consistent with the effects being mediated by changes in striatal circuits, we also observe a preservation of the density of mature spines and length of dendritic arbors in dSPNs in both transgenic transiently elevated dopamine (TED) models, the VGLUT3 KO (cTED) and the eDREADD inducible (iTED) mice. The behavioral and morphological effects are also striatal D1 receptor dependent, as antagonists administered in the striatum of VGLUT3 KO mice before and during striatal DA depletion, prevent the preservation of dSPN mature spines and dendritic arbors as well as normal motor behavior. Recordings of the major excitatory inputs from primary motor cortex to SPNs revealed a profound loss of input strength to both dSPNs and iSPNs following DA depletion in both cTED mice and control wildtype (WT) mice. In contrast, thalamostriatal input strength from PF neurons was not reduced following DA depletion in either genotype. Neither corticostriatal nor thalamostriatal input shifted to favor dSPN over iSPN in the cTED mice. However, paired pulse facilitation of the thalamostriatal inputs to dSPNs and iSPNs, pronounced in both genotypes, was attenuated in depleted WT mice, but remained unaffected in cTED mice. The highly correlated preservation of dSPN morphology and maintenance of paired-pulse facilitation at PF thalamostriatal inputs with normal motor behavior suggests candidates for further mechanistic exploration in these models.

## Materials and Methods

To improve replicability, a list of Key Resources including sources is provided in Table S1.

### Experimental Animal and subject details

Animals were housed in micro-isolator cages on a reverse 12h night/day cycle (10 A.M. lights off, 10 P.M. lights on). All animals had *ad libitum* access to food and water and were treated in compliance with Institutional Animal Care and Use Committee for the University of Pittsburgh. All efforts were made to minimize the number of animals used and to avoid pain and discomfort. In each experiment, approximately equal numbers of males and females were used. *Slc17a8*^−/−^ mice (also referred to as VGLUT3^−/−^ or VGLUT3 KO mice) were backcrossed at least 10 generations to C57BL/6. DAT^cre^ (RRID: IMSR_JAX:020080) and D1^dt^ (RRID: IMSR_JAX:016204) were obtained from Jackson laboratories. For eDREADD experiments, DAT^cre^ mice were used. For electrophysiological eDREADD experiments, DAT^cre^ and D1^dt^ mice were crossed to obtain DAT^cre^; D1^dt^ mice. When testing the duration of normal motor behavior in VGLUT3 KO mice following DA depletion, experiments began between the age of 3-5 months and ended at 8-10 months. For all remaining experiments, mice were 3-6 months in age.

### Animal Groups and Experimental Approach

Experiment 1: SPN spine morphology in a DA depleted state. VGLUT3 KO and WT mice were used. Experiment 2: Effects of transient elevated DA on motor behavior, spine dynamics, and electrophysiology of SPNs in normal and DA depleted states. DAT^cre^; D1^dt^ mice were used for electrophysiology experiments and DAT^cre^ mice were used for the remaining experiments. Mice in these experiments were randomly divided into three cohorts: 1) mCherry+CNO, 2) eDREADD+Saline, or 3) eDREADD+CNO. At 3 months of age, mice were injected with the eDREADD or mCherry virus bilaterally into the SNpc (details below). At least three weeks following viral injection, eDREADD and mCherry animals were given either saline or CNO one hour before the waking cycle (Zeitgeber = 12) for 5 days. These animals then underwent baseline behavioral testing. The injections continued daily for 7 days following DA depletion via 6-OHDA. Animals were assessed behaviorally, one-week following depletion. Animals were euthanized during either the sleeping or waking cycle and assessed for spine and dendrite morphology and electrophysiology.

### Behavior

Light conditions and temperature were kept at ambient levels (∼32 lux, 75°C). For homecage locomotor activity, mice were placed in rat micro-isolator cages inside a photobeam monitoring system (Kinder Scientific) for 72 hours with food and water *ad libitum*. For the *cylinder test*, mice were placed in a clear glass cylinder ∼8 cm in diameter with a clear acrylic sheet on top. The video camera was placed overhead and recorded one 5-minute video per animal per condition. The number of full weight-bearing forepaw contacts (identified as the majority of the forepaw coming in contact with the cylinder) against the cylinder wall was recorded and expressed as total percentage of contacts. Paw dominance is equal to or greater than 70% ipsilateral paw contacts in the cylinder test. Animals were tested 3-4 hours before and after the start of the waking cycle on separate days. Videos with less than 7 total rears were excluded from analysis. All videos were scored by two trained observers unaware of the treatment groups. Their scores were compiled and averaged together.

### Stereotaxic Surgery

All surgical procedures were performed using aseptic technique. Mice were anesthetized with a 3% induction/1-3% maintenance of isoflurane/100% oxygen mixture. Specific details of surgery can be found in (Divito, Steece-Collier et al. 2015). Briefly, mice were first given bilateral injections of either 1 µL of AAV8-hSyn-DIO-hM3D(Gq)-mCherry (RRID: Addgene 44361) or AAV8-hSyn-DIO-mCherry (RRID: Addgene 50459) into the SNpC/VTA (in mm) 3.29 AP, ±1.30 ML, and −4.40 DV. One month later, mice received unilateral partial dorsal striatal lesions with 6-hydroxydopamine hydrochloride (6-OHDA) to assess Parkinsonian motor behavior. Stereotaxic coordinates used for unilateral injection of 6-OHDA (6 µg, Sigma) or saline were (in mm) +0.85 AP, −1.75 ML, and −2.75 DV. During the same surgical session, mice designated for spine analysis received stereotaxic injections of CTB-488 into the SNr (−3.28 AP, ±1.25 ML, and −4.60 DV), whereas all other animals received equivalent injections of saline in place of CTB-488. Injections were administered at 0.25 µL/min with three-minute incubation periods following needle placement and solution injection. Immediately following surgery, mice were sutured and administered ketoprofen (Ketofen 5 mg/kg, Zoetis) and allowed to recover for one week.

Mice also received IP injections of 0.9% saline (0.5 mL) 1-2x daily for 6 days after surgery. Mouse weight was monitored and any that lost >20% of their presurgical weight were excluded. No animals were lost or euthanized after surgery due to surgical complications. 6-OHDA injected animals with less than 70% decrease in TH immunostaining across the rostral-caudal extent of the dorsal striatum were excluded from analysis. Mice where depletion failed does not produce ipsilateral paw dominance in WT mice, justifying exclusion from behavioral analysis. Baseline behavior tests were performed at least three-weeks following eDREADD injections and post-depleted behaviors were assessed for one-week following 6-OHDA depletion. Brains were either harvested for spine analysis or electrophysiology (see below).

For electrophysiology experiments mapping input from specific cortical cell types to striatum, mice from two GENSAT BAC Cre-recombinase driver lines (Tlx3_PL56 for IT-type neurons and Sim1_KJ18 for PT-type neurons) were used (Gerfen, Paletzki et al. 2013). To maximize time for expression, offspring were injected perinatally at postnatal day (P) 0-3 with a Cre-dependent AAV expressing the red-shifted opsin ReaChR (AAV-flex-ReaChR-citrine, Addgene 50955 (Lin, Knutsen et al. 2013)). Pups were cryo-anesthetized with ice for 6-8 minutes and placed in a clay mold stored in a −20 C freezer between uses. The mold aided in positioning pups during injections while maintaining low temperature. Because bregma is not visible at this age, the confluence of the sinuses, approximately corresponding to lambda, was used for alignment. Because the borosilicate glass used for the injection can pass through the skin and skull at this age, and incision is not necessary. The injection was made at 1800 µm anterior and 1100 µm lateral at two depths measured from the skin (850 µm and 650 µm). 100 nL was injected at each site. For thalamic injections at PF, these injections were targeted along two tracts, the first at 1250 µm posterior to bregma, 1000 µm lateral, and at two depths measured from the pia (3200/3500 µm) and the second at 1750 µm posterior to bregma, 500 µm lateral, also at two depths measured from the pia (3200/3500 µm).

### Cannulation Surgery and Infusion Procedures

Mice were anesthetized with a 3% induction/1-3% maintenance of isoflurane/100% oxygen mixture. An incision was made in the midline of the skull and 2 small burr holes were drilled through the skull at stereotaxic coordinates +0.85 AP and −/+1.75 ML. Low profile 26 Ga, 1.75 mm guide cannulas (Plastics One 42593), containing a 33 Ga, 2.75 mm internal dummy cannula (Plastics One 42597) were implanted into these coordinates at a depth of −2.75 DV and secured with GC FujiCem 2 glass ionomer cement (Fisher, 5081153). The wound was sutured closed if any remaining bone or musculature were exposed. Ketoprofen (Ketofen 5 mg/kg, Zoetis) was administered immediately following surgery and the mice allowed to recover for one week. Mice also received IP injections of 0.9% saline (0.5mL) 1-2x daily for 6 days after surgery. Mice were singly housed following surgical procedures to avoid accidental loss of cannulas. Mouse weight was monitored and any that lost >20% of their presurgical weight were excluded. To infuse saline or D1R antagonist (Tocris, SCH23390) bilaterally into the striatum, mice were first pinned down with a cloth, had their internal dummy cannulas removed, and placed the internal infusion cannula (Plastics One, 42595) into the right and left guide cannula. 1 µL of saline or D1R antagonist (SCH23390, Tocris) were infused at a rate of 0.25 µl/min into each hemisphere while the mice moved freely. Three minutes following infusion, the mouse was again pinned down, the infusion cannulas removed, the internal dummy cannula inserted, and the mouse was placed back into the home cage. Prior to behavioral testing, all mice were acclimated to infusion procedures with 3 days of saline injections. D1R antagonist or saline was infused for 7 days prior to 6-OHDA depletion and 7 days after (Figs. S1 and S3).

### DiI labeling and Immunohistochemistry

To analyze spine morphology, mice were euthanized with an overdose of ketamine/xylazine and intracardially perfused with 25 mL PBS (in mm): 7.95 NaCl, 0.20 KCl, 1.425 Na2HPO4, 0.27 KH2PO4, followed by 50mL of cold PBS with 2.5% paraformaldehyde in PBS (para). Brains were dissected and postfixed for one hour in 2.5% para. Brains were then sectioned at 100µm on a vibratome (LEICA VT1000S) and every other section was wet mounted onto a slide. Small DiI

(Thermofisher, D3911) crystals were then placed on the dorsal striatum via glass micropipette. Sections were incubated in PBS at 4°C for two days and subsequently fixed with 4% para for one hour at room temperature. The slides were then coverslipped and stored at 4°C until imaged on a confocal. The remaining slices were fixed in 4% para before being stained for TH immunoreactivity. Slices were washed (3×5 minutes) in PBS and incubated in 1:1000 Rabbit αtyrosine hydroxylase (RRID: AB_390204) containing 0.4%Triton X-100 and 5% normal donkey serum in PBS for 48 hours at room temperature. After washing, these slices were then incubated in 1:500 Alexa Fluor 488 conjugated donkey α-rabbit (RRID: AB_2313584) containing 0.4%Triton X-100 for two hours at room temperature and then mounted with DAPI Fluromount-G (southern Biotech, 0100-20). For electrophysiology experiments, a 1:500 Alexa Fluor 647 streptavidin (RRID: AB_2341101) was added to the secondary solution for neurobiotin labeling. Labeled neurons were imaged on a confocal at 40x with NIS-elements (RRID: SCR_014329) software, then traced and analyzed for Sholl intersections using Simple Neurite Tracer in Fiji-ImageJ (RRID: SCR_002285) software.

### Electrophysiology

Coronal sections (300 µm thickness) were prepared from brains of mice 3-6 months in age of both sexes and euthanized before the onset of the sleeping cycle (10 am). Slices were cut in ice cold, carbogenated, N-methyl-D-glucamine-HEPES solution containing the following (in mM): 93 N-methyl-D-glucamine, 2.5 KCl, 1.2 NaH2PO4, 30 NaHCO3, 20 HEPES, 25 glucose,10 MgSO4, 0.5 CaCl2, 5 sodium ascorbate, 2 thiourea, and 3 sodium pyruvate, pH 7.3. Slices were allowed to recover for 15 minutes at 33°C in this solution before being held at room temperature for 1 hour in carbogenated artificial cerebral spinal fluid (ACSF) as follows (in mM): 125 NaCl, 26 NaHCO3, 1.25 NaH2PO4, 2.5 KCl, 12.5 glucose, 1 MgCl2, and 2 CaCl2. Recordings were made at room temperature using oxygenated ASCF. Whole-cell patch-clamp recordings were performed using borosilicate pipettes (4 − 6 MΩ resistance) filled with an internal solution containing potassium gluconate (in mM): K-gluconate 135, NaCl 1, MgCl2 2, CaCl2 1, HEPES 5, EGTA 5, Mg-ATP 2 and Na4-GTP 0.2, pH 7.3 (∼300 mOsm/L). Cells in which series resistance changed >20% over the course of the recording were excluded from analysis. Experiments were typically performed at −70 mV and for some cases at +40 mV. Data was collected at 10 kHz using an Axopatch 700B (Molecular Devices) and Wavesurfer software (https://wavesurfer.janelia.org/) and analyzed using either Wavesurfer analysis software or with custom routines written in MATLAB (MathWorks). For sCRACM laser scanning photostimulation experiments in corticostriatal recordings (Petreanu, Huber et al. 2007, Petreanu, Mao et al. 2009), data were acquired on a custom-built laser scanning photostimulation microscope (Shepherd, Pologruto et al. 2003). During sCRACM mapping, neurons were held at −70 mV. A blue laser (473 nm, CrystaLaser) was controlled via scan mirrors (Cambridge Technology). Light pulses were controlled with a shutter (4 ms open time) in series with an acousto-optic modulator (1 ms pulse, QuantaTech) to deliver ∼0.5–2 mW at the specimen plane through a low-power objective (UPlanApo 4×, 0.16 NA, Olympus). Laser power was constant during a given day.

Sweeps consisted of 100 ms baseline and 300 ms following onset of the stimulus. The map grid (16 × 16 sites at 50 µm spacing) was centered horizontally over the soma of the recorded neuron and covered the entire SPN dendritic arbor. Maps were repeated 2–4 times and averaged across trials. Evoked current amplitudes were summed to calculate total synaptic input (in nA). LED stimuli for thalamostriatal experiments, including short-term plasticity experiments, were delivered using flashes of a 470 nm LED (Cairn). Sweeps were interleaved and repeated 5 times per stimulus frequency. The maximum current amplitude obtained for cells held at −70 mV was used as a measure of AMPAR mediated values. To estimate the NMDAR currents, we used the current values 65 ms post peak current for cells held at +40 mV. Firing patterns during current clamp recordings were quantified as follows: In current-clamp mode, firing patterns were determined in response to a series of 12 depolarizing current injections (0.8 s duration) that were delivered every 7 s in increasing steps of 25 pA. Firing patterns were routinely elicited from resting membrane potential (RMP). We calculated number of action potentials and average spike frequency. The average firing frequency (f) was calculated from the reciprocal of the interspike interval (ISI) (Prescott and De Koninck 2002).

### Statistics

A summary of all analyses is listed in Table S2. The Kruskal-Wallis Test was used to analyze the percent decrease in TH immunoreactivity in the dorsal striatum. A mixed effect model with repeated measures was used to analyze depleted animals over the course of 20 weeks. For the remaining analyses, we used a two-way ANOVA, repeated measures analysis (*condition vs genotype or condition vs mouse type*). If no significant interaction was found, we reported the main effect. If a significant interaction was found, we used *post-hoc* comparisons with a Bonferroni correction. All data were measured for normality against a Gaussian distribution, using both Shapiro-Wilk and Kolmogorov-Smirnov tests (alpha=0.05). For electrophysiological results we either used a one-way ANOVA with *post-hoc* Tukey’s multiple group comparisons or student’s t-test. Significance was determined at p<0.05. All data are represented as mean ± SEM. All analyses were performed in GraphPad Prism 8 (RRID: SCR_002798) software. *p<0.05, **p<0.01, ***p<0.001, ****p<0.0001

### Resource Availability

Further information and requests for reagents and resources should be directed and will be fulfilled by Rebecca Seal or Bryan Hooks. Raw data obtained for this study are available from the lead contact upon request. Code generated for data analysis is available upon request. Any additional information required to reanalyze the data reported here is available from the lead contact upon request.

## RESULTS

### Transient Elevation of Dopamine (TED) protects motor behavior for weeks after dopamine depletion

We previously reported that striatal DA levels are transiently elevated during the waking cycle, but not the sleeping cycle, in VGLUT3 KO mice, resulting in a corresponding hyperlocomotion (Divito, Steece-Collier et al. 2015). To assess the potential significance of the enhanced dopamine release and motor behavior (also reported in (Gras, Amilhon et al. 2008)), we examined in parallel, circuit and behavioral changes in these mice in a model of Parkinson’s disease. We found that striatal DA enhancement promotes continued voluntary motor control in cTED mice despite ≥ 70% DA loss in the dorsal striatum. Specifically, we tested for motor deficits following unilateral DA depletion induced by intrastriatal injection of 6-OHDA (6 µg) (Brundin, Isacson et al. 1986), assessing the mice for the emergence of ipsilateral paw dominance (significantly greater ipsilateral paw reaches than at baseline) as well as spontaneous contralateral rotations (Divito, Steece-Collier et al. 2015) (Fig. 1B-D). While WT mice exhibited ipsilateral paw dominance one week following intrastriatal 6-OHDA injection, cTED mice did not develop a significant ipsilateral paw dominance until ∼20 weeks post-depletion (Fig. 1D). Analysis of the behavior after the first week (Fig. 1E) showed that cTED mice exhibited no paw preference during either the sleep or wake phase, in contrast to WT mice. Notably, this contrasts with the pre-depletion locomotion differences, which occur only during the waking cycle. The difference in motor behavior between cTED and WT mice was not due to differences in 6-OHDA lesion efficacy, as both groups showed a similar loss of tyrosine hydroxylase (TH) immunoreactivity in the dorsal striatum (∼75%) (Fig. 1F).

**Figure 1.**
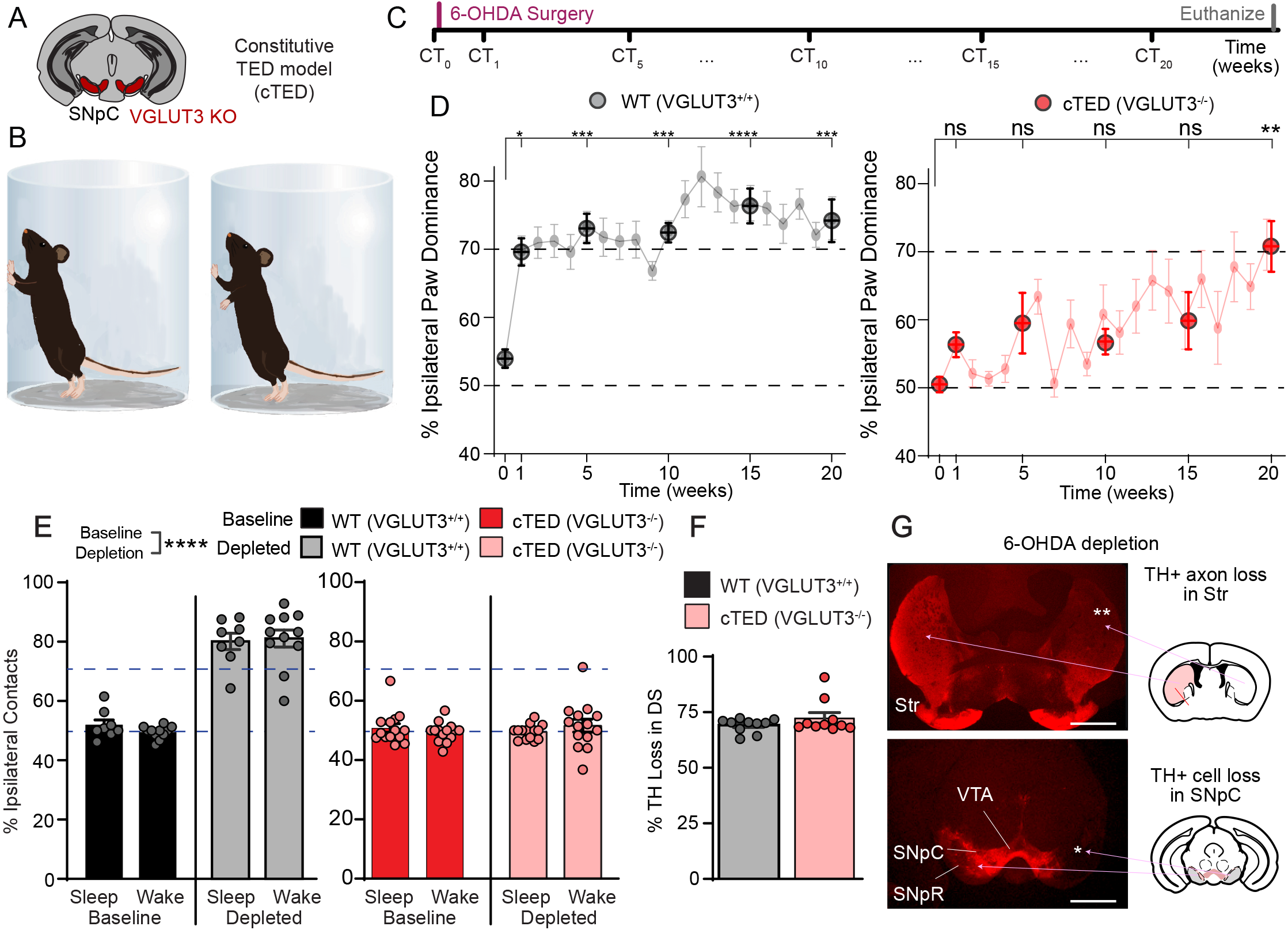
Transient Elevation of DA (TED) affects motor behavior and protects against DA depletion. (A) The constitutive TED model (cTED) uses transgenic VGLUT3 KO mice, which have hyperactivity in the DA system (including SNpC, shown). (B) Cylinder test (CT) experimental paradigm for measuring motor behavior as paw preference following unilateral DA depletion. (C) Timeline of unilateral dorsal striatal 6-OHDA (6 µg) injection for DA depletion (magenta line) and subsequent motor assessment with CT. (D) Control (cTED mice, black) and knockout (cTED, red) mice show significant differences in ipsilateral paw dominance post-depletion. WT mice (black) show ipsilateral dominance at 1 week and after. Dotted lines indicate 50% (normal) and 70% (Parkinsonian) ipsilateral paw reaches, respectively. One-way repeated measures ANOVA with Bonferroni post hoc comparison to baseline *p<0.05, **p<0.01, ***p<0.001, ****p<0.0001, ns = not significant. N=10-13 WT mice at each time point; N=8-10 cTED mice at each time point. (E) Change in ipsilateral contacts one week following depletion. N=8 WT sleeping and N=10 WT waking. N=11 cTED sleeping and N=13 cTED waking. (F) 6-OHDA depletions result in similar reduction in TH+ axons in striatum for both WT and cTED mice (N=10/group), as quantified by percent of striatal area lost (G). Scale bars, 1 mm.

### Enhanced activation of Dopamine Receptor D1 is required for the preservation of motor behavior after dopamine depletion

Because the motor protection in the cTED model could be due to other circuit changes (e.g. in any VGLUT3-associated circuitry), we tested whether this effect on motor behavior involved the hyperactive changes in the DA system. First, we tested whether the neuroprotective effects of the cTED model were mediated by striatal D1 receptors by infusing a D1R dopamine receptor antagonist (SCH23390) into the striatum at the onset of the waking cycle prior to and during 6-OHDA mediated DA depletion. As previously observed, saline-treated cTED mice exhibited increased motor activity in the photobeam monitoring system during the waking cycle and this effect was blocked by treatment with the D1R antagonist (Fig. S1C). One week after DA depletion, D1R antagonist-treated cTED mice showed ipsilateral paw dominance comparable WT control mice, while saline control mice showed bilateral paw reaching behavior typical of cTED mice. These results indicate that striatal D1R-mediated signaling is required for the preserved motor behavior of cTED mice.

### An inducible experimental model recapitulates the dopamine-related phenotypes of constitutive TED mice before and after depletion

This observation supports our hypothesis that normal motor behavior observed in cTED mice after 6OHDA induced depletion results from increased striatal DA levels and enhanced activation of striatal D1 receptors. However, because VGLUT3 is deleted globally (Seal, Akil et al. 2008) in cTED mice, additional phenotypes may contribute to the normal motor behavior. We therefore generated an inducible mouse model designed to recapitulate the elevated DA release and hyperactivity phenotypes observed in cTED mice (Fig. 2A-C). Specifically, we expressed the excitatory Designer Receptor Exclusively Activated by Designer Drugs (eDREADD), hM3Dq (Armbruster, Li et al. 2007, Krashes, Koda et al. 2011) in dopaminergic neurons of the SNpC/VTA (Ventral Tegmental Area) by using DAT^Cre^ mice (Zhuang, Masson et al. 2005). One week later, we injected CTB-488 into the SNpR to label dSPNs. Three weeks later, the eDREADD ligand, clozapine-n-oxide (CNO, 0.1 mg/kg, i.p.) was injected one hour prior to the onset of the waking cycle for 14 consecutive days. The CNO dose was selected to match the locomotor activity of eDREADD mice to that observed in cTED mice during the waking cycle (time of hyperactivity) (Fig. 2E) (Divito, Steece-Collier et al. 2015). On Day 7 of the CNO treatment, we injected 6-OHDA into the dorsal striatum. We refer to transgenic model with eDREADD+CNO as iTED. Two different control cohorts were also included in these analyses: DAT^Cre^ mice expressing eDREADD, injected with saline (eDREADD+Saline, light blue) and DAT^Cre^ mice expressing the mCherry reporter, injected with CNO (mCherry+CNO, dark blue). Collectively, these two conditions are the controls for iTED mice.

**Figure 2.**
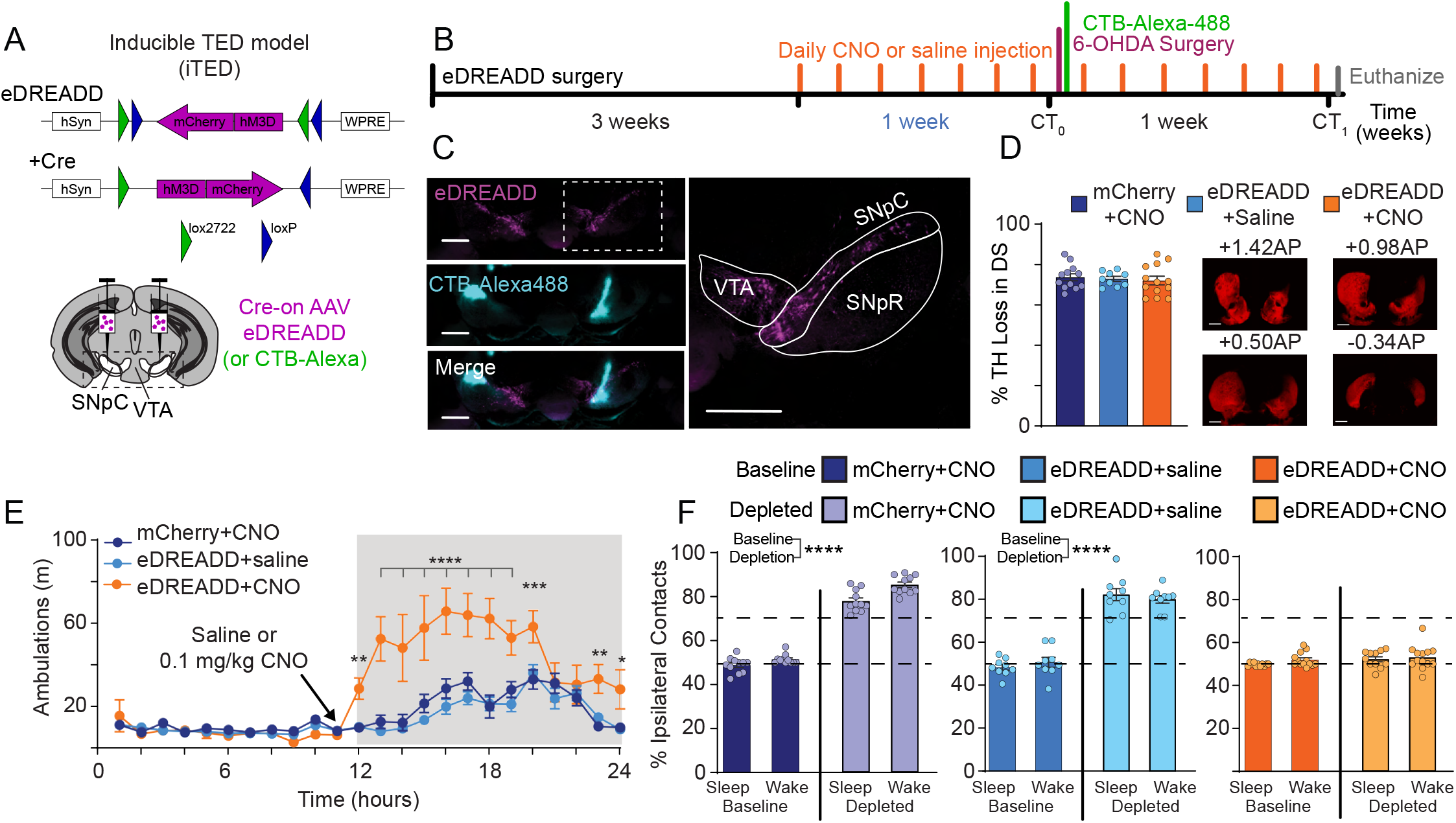
Transient Elevation of DA (TED) affects motor behavior and protects against DA depletion. (A) The inducible TED model in which DAT-Cre mice express eDREADD (hM3Dq) via stereotaxic AAV injection into SNpC. (B) Timeline of Inducible TED activation via eDREADD transfection, daily CNO injection, and DA depletion via intrastriatal 6-OHDA injection. CTB-Alexa-488 is also injected to SNpR to label dSPNs. (C) eDREADD expression in DAT-Cre+ neurons in VTA and SNpC. Left, images hM3Dq-mCherry (magenta) and CTB-Alexa-488 (cyan) of VTA and SNpC (dotted box from (A)). Right, high magnification of VTA (dotted box from (C, left)). Scale bars, 0.5 mm. (D) 6-OHDA depletions result in similar reduction in TH+ axons in striatum for control and eDREADD mice, as quantified by percent of striatal area lost. Scale bars, 0.5 mm. N=11-14/group. (E) CNO injection at the start of the waking cycle results in increased motor activity in eDREADD mice, but not in controls (saline treated eDREADD mice or CNO treated mCherry mice) N=7-10 mice per group. (F) Control (mCherry+CNO, blue, N=12 mice, and eDREADD+saline, cyan, N=11 mice) and eDREADD+CNO (orange, N=14 mice) mice show significant differences in ipsilateral paw dominance post-depletion. Dotted lines indicate 50% (normal) and 70% (Parkinsonian) ipsilateral paw reaches, respectively. One-way repeated measures ANOVA with Bonferroni post hoc comparison to baseline *p<0.05, **p<0.01, ***p<0.001, ****p<0.0001, ns = not significant.

We first tested whether the iTED experimental paradigm preserves contralateral forepaw reaching in the cylinder test following DA depletion. As before, cylinder paw contacts were measured during both sleeping and waking cycles. Before depletion, all cohorts exhibited bilateral paw reaching at both time points. One week after DA depletion, both control cohorts showed ipsilateral paw dominance at both time points (Fig. 2F, p<0.0001). In contrast, iTED mice showed no paw preference in the cylinder test at either time point (Fig. 2F, n.s.). Notably, CNO was not injected on the day of behavioral testing, and the extent of dorsal striatal DA depletion did not differ among cohorts (Fig. 2D). An advantage of the iTED model versus the cTED model is the ability to control the timing of elevated striatal DA levels. We therefore determined the temporal requirements of the eDREADD enhanced DA required to preserve motor behavior. The CNO treatment schedule for the days prior to 6-OHDA treatment was varied (1, 3, or 7 days prior to 6-OHDA injection and 7 days following; Fig. S2). DA loss induced ipsilateral paw preference when measured one week following 6-OHDA using the 1 day schedule. In contrast, the 3 and 7 day schedules showed the preservation of bilateral paw reaches (Fig. S2C). This experiment suggests that induction of the plasticity that protects against motor deficits normally induced by DA loss requires that DA release is enhanced more than one day prior to the 6-OHDA injection.

### Enhanced striatal dopamine release specifically preserves dSPN morphology after dopamine depletion

Our previous work (Divito, Steece-Collier et al. 2015) demonstrated that immature spine density is increased during the waking cycle in a subset of SPNs in cTED mice. We next examined whether the subset of SPNs showing increased immature spines belongs to a specific SPN subtype. To label the dSPNs specifically, we injected CTB-488 into the SNpR. One week later, mice were sacrificed during the waking and sleeping cycles and striatal slices prepared. The spine analyses were carried out in the same animals used for behavior. DiI was sparsely added to the acute slices to label SPN membranes, which were then analyzed for SPN spine density and morphology on secondary dendrites. We subdivided spines by morphology assigning them as mature spines (mushroom, Fig. 3) or immature spines (either long/thin or stubby; Fig. S3). These analyses showed that the increased immature spines of cTED mice, compared to WT control mice, during the waking cycle are on only direct pathway SPNs (WT, black) vs cTED mice (red, Fig. S3A; p<0.0001). In contrast, immature spines on dSPNs during the sleeping cycle are not increased. iSPNs spine density during both sleeping and waking time points is the same in WT and cTED mice (Fig. S3A).

**Figure 3.**
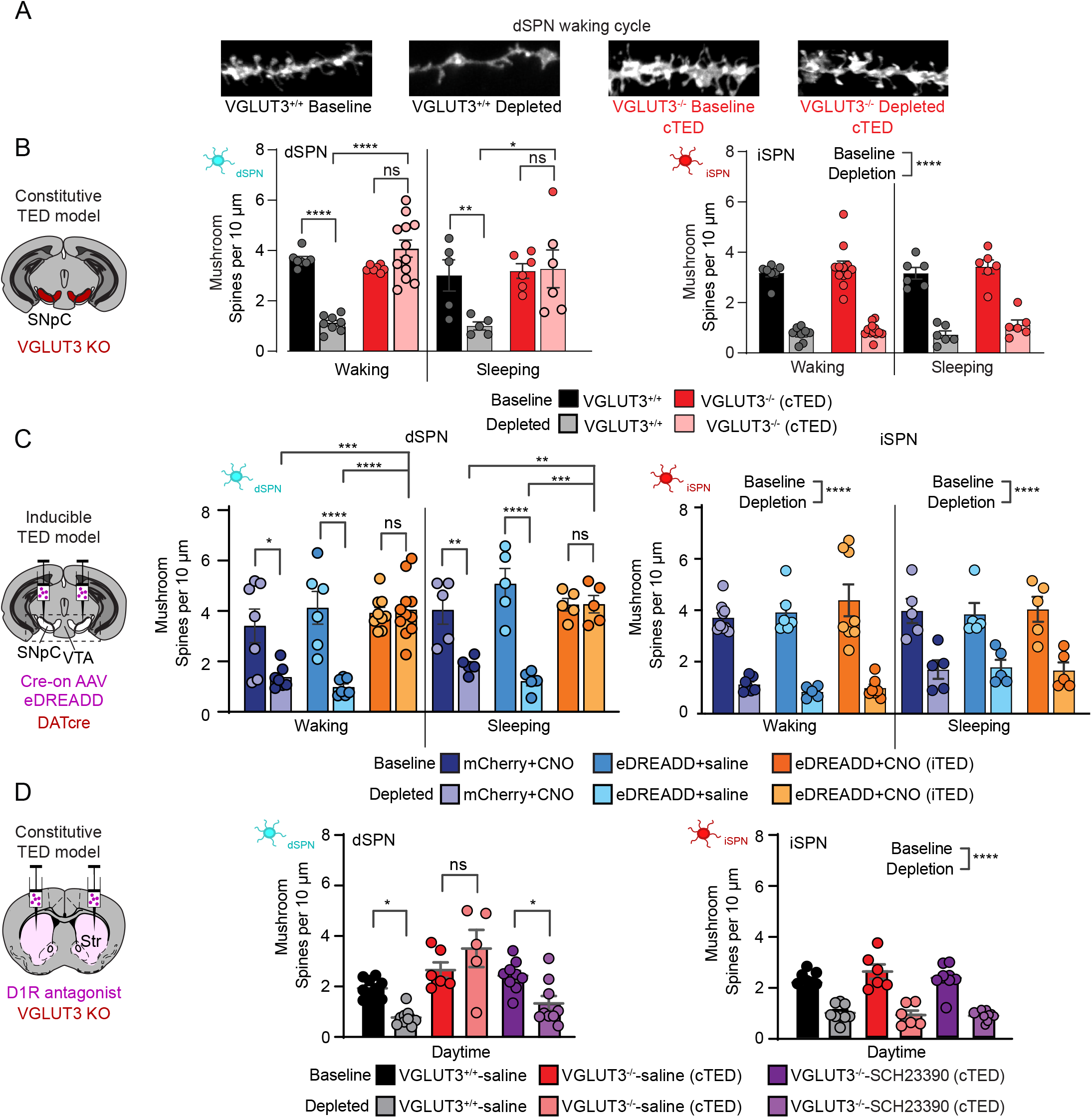
Transient Elevation of DA (TED) preserves mature dSPN spines but not iSPN spines following DA depletion. (A) Representative images of DiI-filled dSPN dendrites during the waking cycle from WT (black) and Constitutive TED (red) mice. (B) Spine data for the constitutive TED model. After DA depletion, mature (mushroom) spine densities on dSPNs are significantly decreased in WT mice (black vs gray) but are preserved in Constitutive TED mice (red vs pink bars) for both waking and sleeping phases across the circadian cycle. Circles represent number of spines per dendrites measured. iSPN spine densities are similar at baseline in Constitutive TED and WT mice and all spines are decreased similarly by depletion in both genotypes. Two-way ANOVA (genotype x condition), no significant interaction, main effect for condition is reported. N=3-4 mice and n=6-12 dendrites/group for all mice (waking and sleeping). (C) Spine data for the inducible TED model. After depletion, mature (mushroom) spine density is preserved in eDREADD+CNO mice (orange) but not in saline-treated control mice (light blue) across the circadian cycle. Significant interaction by 2-way ANOVA (cohort x condition) for mushroom spines. Bonferroni multiple comparisons. Significant main effect for condition. Depletion decreased iSPN spine densities to the same extent in eDREADD+CNO mice (orange) and controls (light and dark blue bars). No interaction by 2-way ANOVA (cohort x condition). Significant main effect for condition. N=3-4 mice and n=7-12 dendrites/group for all mice (waking and sleeping). (D) Spine data for the D1R antagonist experiment. At baseline, dSPN spine densities were similar between cohorts. After DA depletion, mature (mushroom) spine densities of Constitutive TED (D1 antagonist infused) were decreased similarly to the WT (saline infused), while control Constitutive TED (saline infused) showed the expected preserved mushroom spine densities. Significant interaction by 2-way ANOVA (cohort x condition) with Bonferroni multiple comparisons. Baseline iSPN spine densities were similar between cohorts. After depletion, spine densities were significantly decreased across cohorts. All measurements taken during the sleeping cycle. No interaction by 2-way ANOVA (cohort x condition). Significant main effects of condition. *p<0.05, **p<0.01, ***p<0.001, ****p<0.0001, ns = not significant. N=3-4 mice and n=6-9 dendrites assessed/group for all groups (saline treated WT mice, saline treated cTED mice, and D1R antagonist treated cTED mice).

This increase in density of immature spines only on dSPNs of cTED mice during the time of elevated DA then prompted us to examine the density of spines following dopamine depletion, with a particular interest in the mature spines, which are found to be reduced in both SPN subtypes in animal models of PD and post-mortem brains of PD patients. An increase in mature spines on dSPNs following DA depletion could preserve synaptic connections specifically to the direct pathway, thus preserving the normal balance of striatal output and motor behavior (Stephens, Mueller et al. 2005, Smith, Raju et al. 2009). The concept of a DA-mediated increase in immature spines that subsequently mature with DA depletion is based on the cocaine literature, where it has been shown that a daily administration of cocaine (which elevates DA) produces immature spines, which then mature following a 1–2 week period of cocaine withdrawal (in this case 6-OHDA mediated DA depletion) (Huang, Lin et al. 2009, Lanciego, Luquin et al. 2012, Lee, Ma et al. 2013).

We thus similarly quantified the density of mature spines in SPNs across conditions and found that unlike in control mice (Fig. 3A-B, p<0.05), the density of mature spines in dSPNs was not reduced in cTED mice one week following DA depletion, when assayed either during the waking or the sleeping cycles (Fig. 3B, left). The results for iSPNs of cTED mice, however showed a loss of mature spines following DA depletion at both time points similar to WT control mice (Fig. 3B, right). We confirmed that the degree of DA depletion measured using TH immunoreactivity was similar between cTED and WT control mice (Fig. 1F). Notably, we also observed that dSPN-specific preservation of mature spine density in cTED mice required enhanced D1R activation, since antagonism of D1R via infusion of the antagonist SCH23390 for 7 days prior and 7 days following 6-OHDA injection resulted in a loss of mature spines only on dSPNs in cTED mice, similar to saline-treated WT control mice (Fig. 3D).

We next examined SPN spine densities in iTED and control cohorts (Figs. 3C and S3B). In non-depleted conditions, mice were injected with 0.1 mg/kg CNO or saline, just prior to the onset of the waking cycle and brains harvested 3 hours later. Spine analyses using acute slices showed an increase in the densities of long/thin immature spines on dSPNs of only the iTED mice, similar to what we observe in cTED mice (Fig. S3B, left; p<0.05 vs. eDREADD+saline and p<0.0001 vs. mCherry+CNO). Also similar to cTED mice, mature spine densities of iTED mice were the same as controls. In dopamine depleted conditions, iTED mice showed greater mature spine densities in dSPNs than controls, similar to the cTED mice (Fig. 3C, left). Also similar to the cTED mice and controls, spine densities on iSPNs were significantly reduced across iTED mice and controls at both time points (Figs. 3C and S3B right, p<0.0001).

Reduced length and complexity of dendritic arbors on both types of SPNs after DA depletion has been consistently reported for rodent models (Day, Wang et al. 2006, Fieblinger, Graves et al. 2014, Gagnon, Petryszyn et al. 2017, Villalba and Smith 2018), the MPTP primate model (Smith and Villalba 2008, Smith, Villalba et al. 2009, Villalba and Smith 2010) and postmortem brains of PD patients (Stephens, Mueller et al. 2005, Zaja-Milatovic, Milatovic et al. 2005). To assess dendritic architecture, we performed Sholl analyses on SPNs under non-depleted and depleted conditions (Fig. 4A,B). Under non-depleted conditions, the degree of dendritic arbor branching by dSPNs of iTED mice did not differ significantly from controls (Fig. 4B, top left). After depletion, however, the extent of branching of dSPNs was significantly greater in iTED mice than controls (Fig. 4B top right). In contrast, the branching of iSPNs did not differ between iTED mice and control before or after depletion (Fig. 4B, bottom). Results of total dendritic length matched what we observed for mature spine densities and dendritic branching. The dSPNs in control cohorts but not iTED mice showed significantly decreased arbor length after DA depletion (mCherry+CNO, baseline vs depleted p<0.05, eDREADD+Saline, baseline vs depleted p<0.01; iTED mice, baseline vs depleted, n.s.; Fig. 4C left). For iSPN, all cohorts showed significantly decreased total dendritic lengths after DA depletion (p<0.0001; Fig. 4C right). Thus, the data mirror findings observed in cTED mice. Specifically, transiently increasing striatal DA release enhances immature spine density on dSPNs under non-depleted conditions while doing so prior to and during depletion instead preserves the mature spine density and the dendritic arbor length and branching on only dSPNs as well as normal motor behavior.

**Figure 4.**
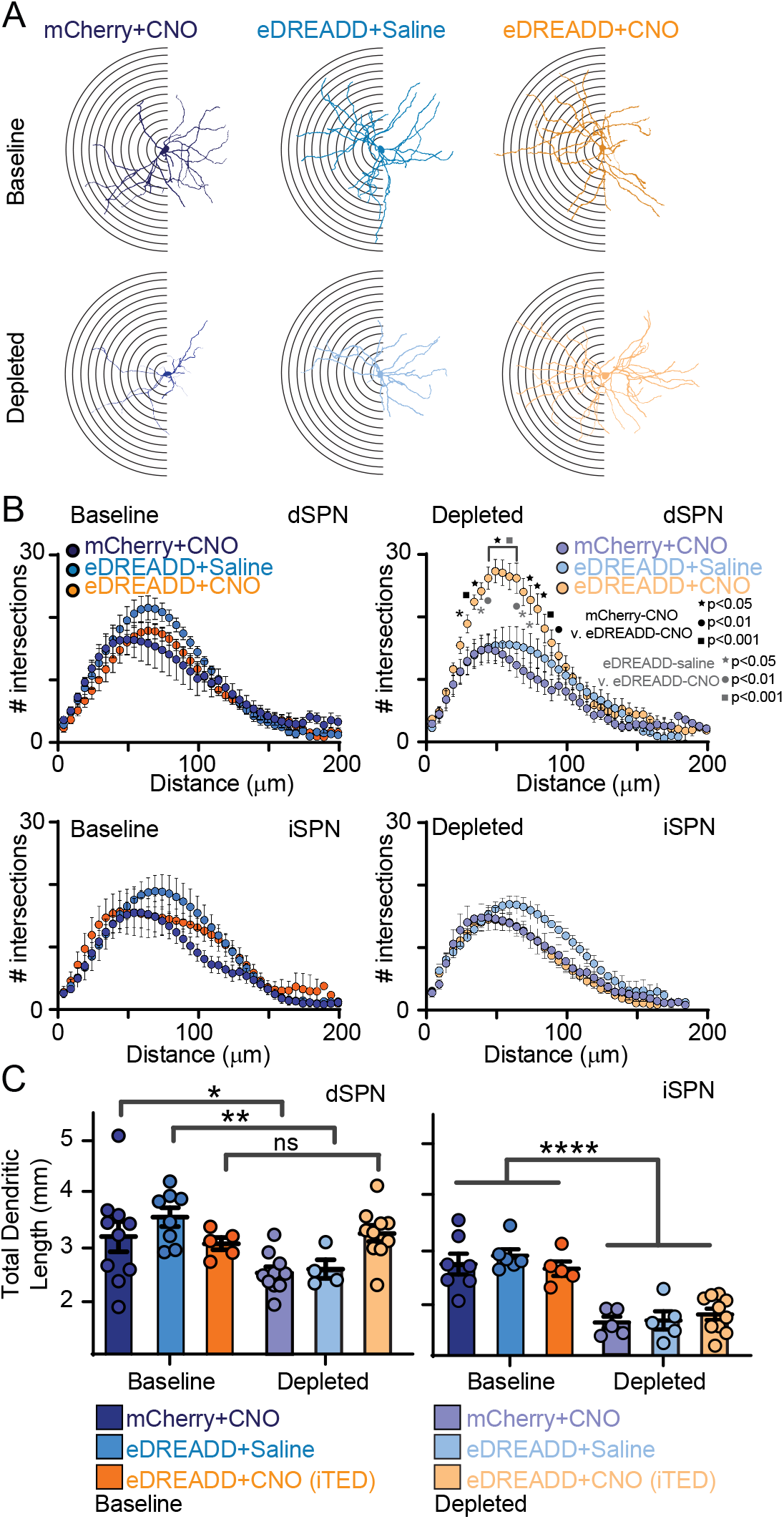
Transient Elevation of DA (TED) preserves mature dSPN dendrites following DA depletion. (A) Scholl analysis of dendritic arbor size in the inducible TED model. Dendritic arbor extent is estimated by the number of times the dendrites of a single filled dSPN or iSPN crosses rings every 5 mm from the soma. (B) Quantification for basline (left) and depleted (right) mice, with dSPN on top and iSPN at bottom. (C) The sum of total dendritic length is affected by DA depletion for dSPNs in control cases (mCherry+CNO and eDREADD+saline) but not inducible TED cases (eDREADD+CNO). iSPNs are affected similarly in all cases. One-way repeated measures ANOVA with Bonferroni post hoc comparison to baseline *p<0.05, **p<0.01, ***p<0.001, ****p<0.0001, ns = not significant. N=3-4 mice/group and n=4-11 dendrites/group.

### Intrinsic electrophysiological properties of SPNs after dorsal striatal dopamine depletion

The intrinsic physiological properties of SPNs influence neuronal excitability and output of SPNs and thus motor deficits in models of PD (Fieblinger, Graves et al. 2014). We therefore characterized the intrinsic properties of SPNs in non-depleted and depleted cTED mice and WT controls. Consistent with previous reports (Fieblinger, Graves et al. 2014), the number of action potentials (AP) and firing frequency (see methods for details of frequency calculation) evoked by current injection measured for dSPNs was increased in DA depleted WT mice vs non-depleted WT mice (Fig. 5D-E left, black and red). The dSPNs in cTED mice following depletion also showed an increase in APs, though to a lesser degree than those of WT mice (Fig. 5D-E left, blue and magenta). In contrast, APs evoked in iSPNs were not affected across groups (Fig. 5D,E, right; n.s.). The resting membrane potentials (Vrest) of dSPNs (Fig. 5F) became more depolarized in WT mice with DA depletion (p<0.01, WT naïve vs depleted). In contrast, Vrest measured in dSPNs of cTED mice was not altered by depletion (n.s. cTED naïve and depleted). The Vrest of dSPNs in non-depleted cTED and WT mice did not differ (n.s. cTED non-depleted and WT non-depleted). Vrest of iSPNs (Fig. 5F) were also more depolarized for depleted mice compared to WT naïve mice (p<0.01, WT naïve vs depleted) but showed a similar decrease in cTED mice (n.s., WT naïve vs depleted). The relative changes can be clearly seen in the plot of the mean number of APs by dSPNs and iSPNs across conditions and genotypes (Fig. 5G). Thus, two measures of increased excitability observed in dSPNs of WT mice with depletion, APs in response to current injection and Vrest, were less pronounced and absent, respectively in cTED mice.

**Figure 5.**
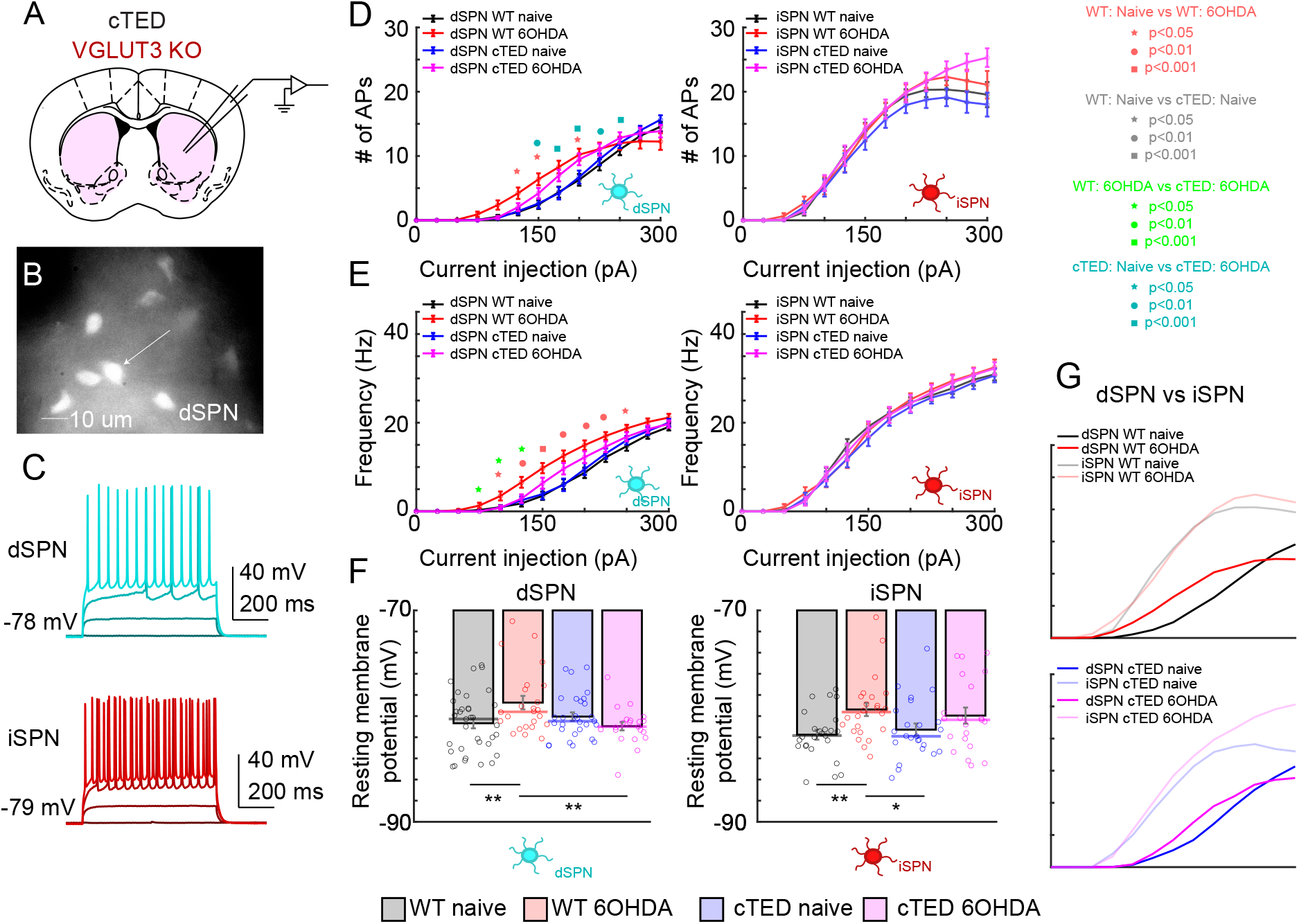
Recording excitability in dSPNs and iSPNs. (A) Recording excitability of dSPN (transgenically labeled with Drd1a-tdTomato) or unlabeled iSPN cells. (B) 60x fluorescence image of tdTomato+ dSPN in a PL56-Cre+;Drd1a-tdTomato+ striatal brain slice. (C) Representative current injection traces from dSPN (blue, top) and iSPN (red, bottom) for selected current injection magnitudes. Numbers on left bottom indicate their RMP. (D) Intrinsic excitability (# AP per current injected) for dSPN (left) and iSPN (right) in control and 6-OHDA depleted mice in WT or VGLUT3 KO background. cTED in legend, cTED. Statistics: One-way ANOVA with Tukey’s post hoc multiple comparison of groups and student’s t-test: *p<0.05, **p<0.01, ***p<0.001, ****p<0.0001, ns = not significant. WT, N=34 neurons; WT depleted, N=24; cTED, N=29; cTED depleted, N=21. (E) Alternate intrinsic excitability via the firing frequency that was evaluated as the average of inverted inter-spike intervals for each current injection steps that elicited at least two APs. Note: Each group has an assigned color, where symbols + error bars show the magnitude of differences. (F) Resting membrane potential. Bar plot, mean. Scatter plot, raw values. Solid line is median. Statistics and sample size the same across (D-F). (G) Comparison of mean #APs, plotted as in (D) to compare dSPN and iSPN excitability.

### Loss of excitatory input from IT- and PT-type cortical projections to dSPNs and iSPNs

Primary motor cortex (M1) provides the main excitatory input to dorsal striatal SPNs. This input is important for both motor learning and execution of movement and thus, was specifically selected for analysis (Currie, Ammer et al. 2022, Park, Phillips et al. 2022, Shinotsuka, Tanaka et al. 2023). Cortical input to striatum originates from two main populations of layer 5 pyramidal neurons (Shepherd 2013, Harris and Shepherd 2015): Intra-Telencephalic (IT-type) and Pyramidal Tract-type (PT-type) neurons (Fig. 6A,B). IT-type cells project to ipsilateral and contralateral striatum and cortex, but not to other subcortical targets, while PT-type neurons project to ipsilateral striatum and numerous subcortical structures as well as spinal cord (Wilson 1987, Shepherd 2013). We therefore tested whether the preserved mature spines and arbors on dSPNs observed in cTED mice after DA depletion reflect the selective preservation of cortical inputs to dSPNs (Fig. 6). Well-characterized Cre driver lines that selectively label IT-type neurons (Tlx3_PL56-Cre) or PT-type neurons (Sim1_KJ18-Cre) (Gerfen, Paletzki et al. 2013, Hooks, Papale et al. 2018) were used for channelrhodopsin-2 Assisted Circuit Mapping (CRACM; (Petreanu, Huber et al. 2007, Petreanu, Mao et al. 2009)). AAV1 encoding Cre-dependent red-shifted opsin, ReaChR, fused to mCitrine (Lin, Knutsen et al. 2013, Hooks, Lin et al. 2015) was injected into the forelimb primary motor cortex (M1) of each Cre line at P0-P3. Acute slices were prepared at P24-158 for recording the excitatory postsynaptic currents (EPSCs) in both dSPNs and iSPNs of the dorsolateral striatum. Drd1-tomato reporter mice were used to identify dSPNs (Ade, Wan et al. 2011). Tomato-negative neurons displaying spiny morphology upon post-hoc analysis were classified as iSPNs. Evoked current amplitudes were summed to calculate total synaptic input (in nA).

**Figure 6.**
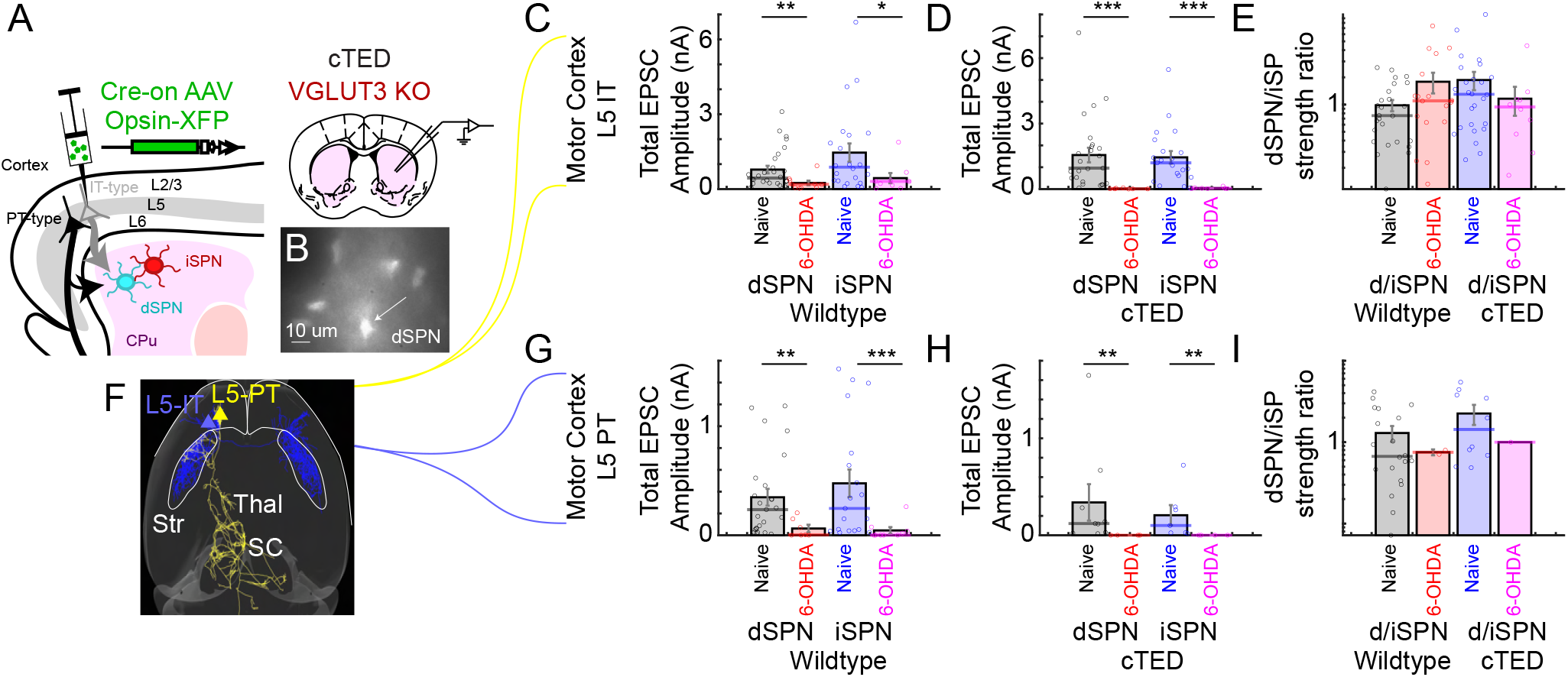
Recording from IT-type and PT-type input to dSPNs and iSPNs. (A) Stereotaxic AAV injection of Cre-dependent opsin makes IT-type (or PT-type, not shown) axons excitable by applied blue light for recording input to dSPN (transgenically labeled with Drd1a-tdTomato) or unlabeled iSPN cells. Recordings test input strength and clustering. (B) 60x fluorescence image of tdTomato+ dSPN in a PL56-Cre+;Drd1a-tdTomato+ striatal brainslice. (C) Total EPSC amplitude to dSPN (left) and iSPN (right) from IT-type neurons in WT mice before or after DA depletion (6-OHDA treatment). *, p<0.05; **, p<0.01; ***, p<0.001, Mann-Whitney U-test. Thick line is mean value. For dSPN, WT N=28 neurons; WT depleted N=10. For iSPN, WT N=22 neurons; WT depleted N=9. (D) Total EPSC amplitude to dSPN (left) and iSPN (right) from IT-type neurons in cTED mice before or after DA depletion (6-OHDA treatment). *, p<0.05; **, p<0.01; ***, p<0.001, Mann-Whitney U-test. Thick line is mean value. For dSPN, cTED N=25 neurons; cTED depleted N=7. For iSPN, cTED N=21 neurons; cTED depleted N=7. (E) dSPN/iSPN strength ratio for IT-type inputs. (F) Schematic showing IT-type and PT-type axons. (G) Total EPSC amplitude to dSPN (left) and iSPN (right) from PT-type neurons in WT mice before or after DA depletion (6-OHDA treatment). *, p<0.05; **, p<0.01; ***, p<0.001, Mann-Whitney U-test. Thick line is mean value. For dSPN, WT N=25 neurons; WT depleted N=7. For iSPN, WT N=19 neurons; WT depleted N=9. (H) Total EPSC amplitude to dSPN (left) and iSPN (right) from PT-type neurons in cTED mice before or after DA depletion (6-OHDA treatment). *, p<0.05; **, p<0.01; ***, p<0.001, Mann-Whitney U-test. Thick line is mean value. For dSPN, cTED N=9 neurons; cTED depleted N=4. For iSPN, cTED N=7 neurons; cTED depleted N=4. (I) dSPN/iSPN strength ratio for PT-type inputs.

Under non-depleted conditions, IT-type projection neurons elicited monosynaptic currents in dSPNs and iSPNs roughly equally in both cTED mice and WT controls (Fig. 6C-E). In WT mice, iSPN input was non-significantly favored compared to dSPN input. In cTED mice, iSPN input was also about the same as dSPN input. Following DA depletion, however, stimulating IT-type inputs in WT mice produced significantly less current in both dSPNs (p=0.0055, naïve vs depleted) and iSPNs (p=0.033, naïve vs depleted), with ∼70% less inputs than naïve mice. Surprisingly, the reduction in EPSCs was much greater in cTED mice for both iSPNs (∼97% loss; p<0.0001, naïve vs depleted) and dSPNs (∼99% loss; p<0.0001, naïve vs depleted). Despite the substantial reduction in synaptic input, the ratio of IT-type input to dSPNs and iSPNs remained largely unchanged following DA depletion in both genotypes (Fig. 6E).

Results were similar for PT-type inputs (Fig. 6G,H). It is notable that the overall strength of PT-type inputs was weaker than IT-type inputs, ∼33% of the strength. In primates, these are believed to be much weaker (Pasquereau, DeLong et al. 2016), though the PT-type corticostriatal projection is still anatomically substantial in mouse (Hooks, Papale et al. 2018). For controls, iSPN input was again non-significantly favored (Fig. 6I) compared to dSPN input. For cTED mice, iSPN input was closer to the same as dSPN input (Fig. 6C). Testing the effects of DA depletion on PT-type afferents, we observed a more severe loss of input than for IT-type inputs. For iSPN input in WT mice, ∼91% of the strength was lost (p=0.0006, naïve vs depleted) and for dSPN input, ∼82% was lost (p=0.0071, naïve vs depleted). The PT-type loss of synaptic connectivity in cTED mice was even more severe, with ∼1% of the original input remaining to both iSPN (p<0.0061, naïve vs depleted) and dSPN (p<0.0028, naïve vs depleted). Thus, DA depletion produced a profound loss of both IT-type and PT-type corticostriatal input to dSPNs and iSPNs in both cTED and WT mice, despite the preservation of mature spines and dendritic arbors of dSPNs in cTED mice. This apparent dissociation between the preserved dSPN morphology and loss of M1-derived synaptic input suggests that the preserved dSPN spines may receive alternative excitatory inputs and/or that preserved dSPN spines do not form functional synapses.

### Maintained excitatory thalamostriatal input

A second major source of glutamatergic input to the dorsolateral striatum originates from motor thalamus, including the SNpR-recipient parafascicular nucleus (PF; (Lee, Wang et al. 2020)). These thalamic nuclei make convergent synaptic contact on the spines and shafts of direct and indirect pathway SPNs (Smith, Raju et al. 2009). Within the same neuron, the NMDAR/AMPAR ratio is greater for excitatory responses evoked from the thalamostriatal pathway than for those evoked from the corticostriatal pathway (Smeal, Keefe et al. 2008). Ideal Cre-driver lines to segregate distinct thalamic nuclei have been challenging to identify, and subtypes of thalamic relay neurons within a given mouse nucleus have not shown substantial genetic variation (Phillips, Schulmann et al. 2019). We therefore mapped the strength of these inputs via CRACM (Petreanu, Huber et al. 2007, Petreanu, Mao et al. 2009)) with the kinetically rapid opsin Chronos (Klapoetke, Murata et al. 2014) in a non-Cre dependent AAV vector (Fig. 7A). The injection site in PF was confirmed by assessment of thalamic brain slices made following sectioning for recording. A representative image of the virally transduced thalamic target is shown in Fig. 7B. The rapid kinetics were also used to evaluate synaptic strength during trains of stimuli (Fig. 8 below).

**Figure 7.**
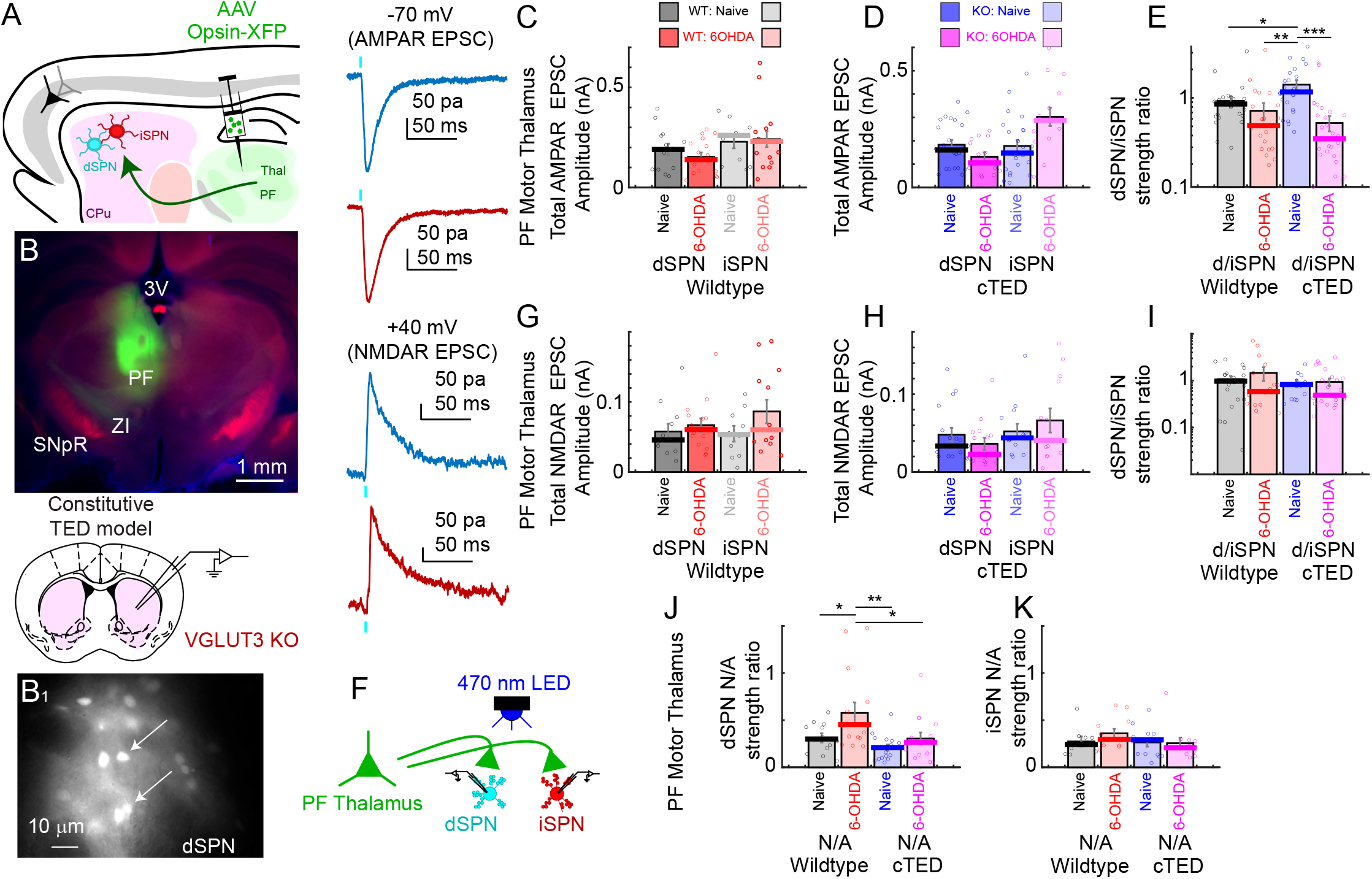
Recording from PF Motor Thalamus input to dSPNs and iSPNs. (A) Stereotaxic AAV injection of Cre-dependent opsin makes PF Motor Thalamus axons excitable by applied blue light for recording input to dSPN (transgenically labeled with Drd1a-tdTomato) or unlabeled iSPN cells. Recordings test input strength and clustering. (B) Fluorescence image of the injection site in PF (green), with 60x fluorescence image (B1) of tdTomato+ dSPN in a PL56-Cre+;Drd1a-tdTomato+ striatal brainslice. (C) Total EPSC amplitude to dSPN (left) and iSPN (right) from PF Thalamic neurons in WT mice before or after DA depletion (6-OHDA treatment). Thick line is mean value. For dSPN, WT N=14 neurons; WT depleted N=17. For iSPN, WT N=11 neurons; WT depleted N=16. (D) Total EPSC amplitude to dSPN (left) and iSPN (right) from PF Thalamic neurons in cTED mice before or after DA depletion (6-OHDA treatment). Thick line is mean value. For dSPN, cTED N=23 neurons; cTED depleted N=15. For iSPN, cTED N=24 neurons; cTED depleted N=15. (E) AMPAR mediated dSPN/iSPN strength ratio for PF Motor Thalamus inputs. *, p<0.05; **, p<0.01; ***, p<0.001, One-way ANOVA with Tukey’s post hoc multiple comparison. (F) Schematic of recording inputs to SPNs. (G) Total NMDAR amplitude to dSPN (left) and iSPN (right) from PF Thalamic neurons in WT mice before or after DA depletion (6-OHDA treatment). Thick line is mean value. For dSPN, WT N=12 neurons; WT depleted N=15. For iSPN, WT N=10 neurons; WT depleted N=13. (H) Total NMDAR amplitude to dSPN (left) and iSPN (right) from PT-type neurons in cTED mice before or after DA depletion (6-OHDA treatment). Thick line is mean value. For dSPN, cTED N=17 neurons; cTED depleted N=15. For iSPN, cTED N=14 neurons; cTED depleted N=13. (I) dSPN/iSPN strength ratio for PF Motor Thalamus inputs for NMDAR inputs. (J,K) AMPAR/NMDAR input ratio for input to dSPN (J) and iSPN (K). *, p<0.05; **, p<0.01; ***, p<0.001, One-way ANOVA with Tukey’s post hoc multiple comparison (J).

**Figure 8.**
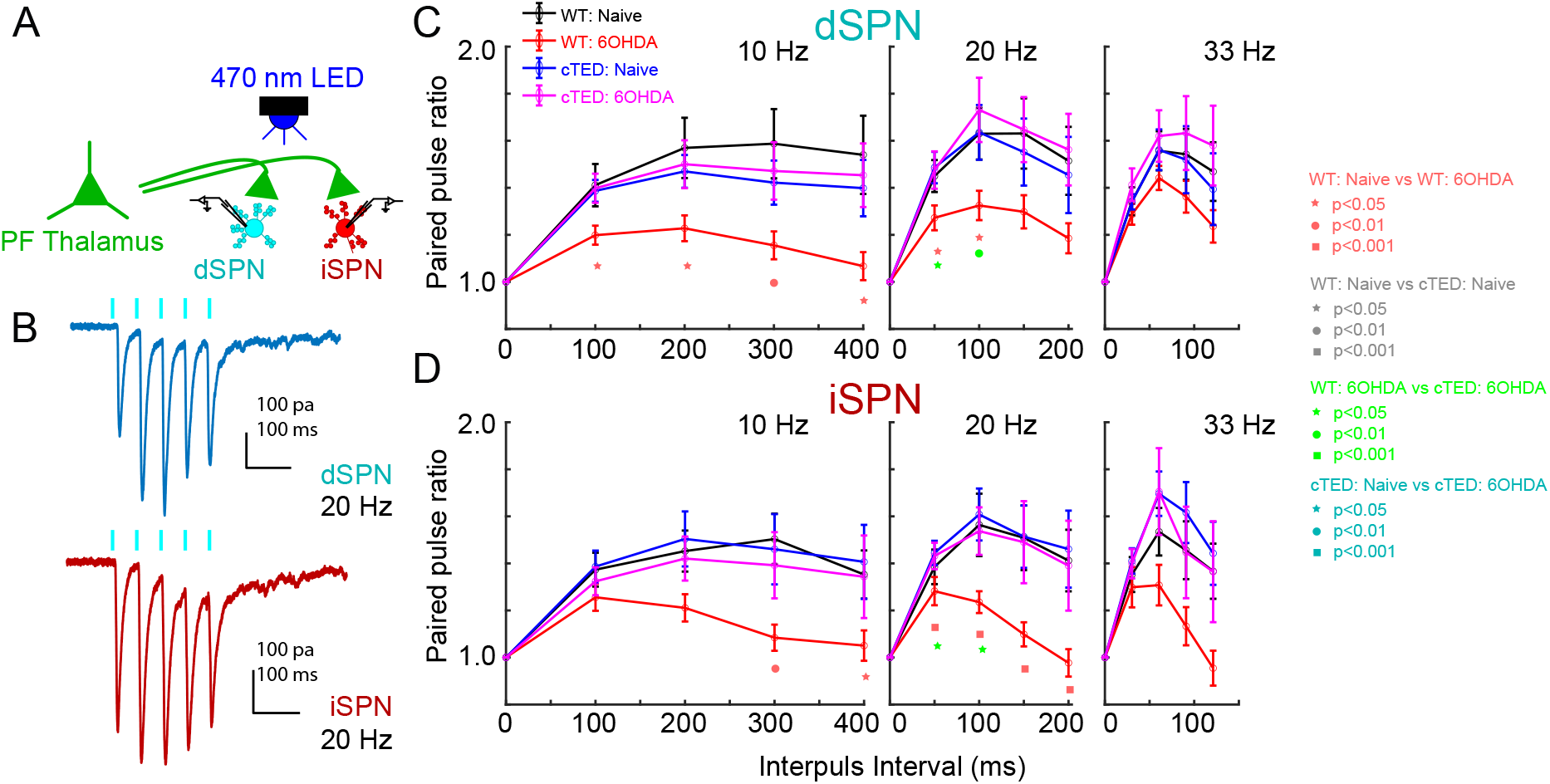
Thalamostriatal short term plasticity following DA loss in control and TED mice. (A) Thalamic AAV injections express Chronos-GFP in PF nuclei. PF injection site verified post hoc. (B) Whole cell recordings in brain slice track EPSC amplitude at −70 mV during pulse trains at 10, 20, and 33 Hz. 20 Hz illustrated for a WT naive slice in dSPN (blue) and iSPN (red) recordings. For dSPN (C) and iSPN (D), short term facilitation is present during 10, 20, and 33 Hz trains for WT and cTED mice in control conditions. After DA depletion by intrastriatal 6-OHDA injection, paired pulse ratio (PPR) is deficient in wildtype mice but preserved in TED mice. For dSPN, WT N=17 neurons; WT depleted N=15; cTED N=14; cTED depleted N=14. For iSPN, WT N=15 neurons; WT depleted N=14; cTED N=17; cTED depleted N=15.

First, we compared the strength of synaptic inputs from PF, recording at −70 mV to measure AMPAR-mediated currents (as in the corticostriatal recordings) as well as at +40 mV to measure NMDAR-mediated currents. As shown in Fig. 7C-D, for both non-depleted WT and cTED mice, AMPAR-mediated thalamic input to iSPNs was similar to the input to dSPNs. The effects of DA depletion on AMPAR-mediated inputs to dSPNs did not differ significantly between WT and cTED mice. We did observe a non-significant increase in the input to iSPNs, and a slight decrease in the input to dSPNs following DA depletion in WT mice (Fig. 7C). Similar effects were observed in the cTED mice, with a slight increase in input to iSPNs and a slight decrease in input to dSPNs (Fig. 7D). Nevertheless, none of the comparisons of the elicited currents described above were significant. To evaluate pathway balance, we calculated dSPN/iSPN ratios of AMPAR-mediated currents (Fig. 7E). Compared to the non-depleted condition, the dSPN/iSPN AMPAR ratio significantly decreased in cTED (0.54 ± 0.10, N = 29; p < 0.0001) but not the WT (0.73 ± 0.15, N = 21; p > 0.05) mice after DA depletion. It is worth noting that before depletion, the ratio was greater for cTED mice (1.38 ± 0.16, N = 24) with respect to WT mice (0.93 ± 0.11, N = 25; p = 0.03).

We also recorded and similarly analyzed the NMDAR-mediated component of PF-elicited glutamatergic currents. In both the cTED and control mice, this component of PF inputs to iSPNs and dSPNs were similar between SPN types and between genotypes (Fig. 7G, H). With respect to the effects of DA depletion, we observed similar NMDAR currents in both iSPNs and dSPNs in control and cTED mice (Fig. 7G,H). Currents were slightly increased in iSPNs and slightly decreased in dSPNs (Fig. 7H) but these effects were non-significant. With respect to the balance of input to dSPNs and iSPNs, WT mice showed dSPN/iSPN ratios of 1.08 ± 0.18 at baseline and 1.46 ± 0.46 after depletion, while cTED mice showed ratios of 0.89 ± 0.15 at baseline and 0.94 ± 0.16 after depletion. Thus, no significant differences in the NMDAR-mediated inputs to SPNs or dSPN/iSPN ratios were observed between genotype or depletion status (Fig. 7I).

NMDAR- and AMPAR-mediated currents were re-examined in the form of NMDAR/AMPAR ratios to assess whether there were changes in receptor composition at functional synapses. Often an increase in NMDAR/AMPAR current ratio implies a greater proportional influx of calcium (Ca²⁺) signaling at a synapse, and strengthening the synaptic signaling (Lisman, Fellous et al. 1998, Popescu, Saghyan et al. 2007). In WT mice, the NMDAR/AMPAR ratio was significantly greater after DA depletion for dSPNs (p = 0.048, baseline vs depleted; Fig 7J) but not for iSPNs (n.s., baseline vs depleted; Fig 7K). In cTED mice, the ratios were similar between depleted and not depleted conditions for both dSPNs and for iSPNs (Fig. 7K). However, the ratios were significantly lower in dSPNs of cTED mice compared to WT after depletion (p = 0.03, WT depleted vs. cTED depleted; Fig. 7J). Thus, these data reveal some changes in the postsynaptic NMDAR composition at thalamostriatal synapses in cTED mice, but presynaptic alterations may also contribute to the preservation of motor function in cTED mice.

We therefore assessed presynaptic release probability changes using trains of pulses to study short term plasticity at the thalamostriatal (PF) input to SPNs in WT and cTED mice following DA depletion (Figure 8). This type of plasticity is especially relevant for thalamostriatal input, since thalamic relay neurons may fire in bursts (McCormick and Pape 1990, Sherman and Guillery 2006). Specifically, we measured the paired-pulse ratio (PPR) elicited by 5 stimuli at 10, 20 and 33 Hz for PF inputs to dSPN and iSPN. To maintain responsivity during rapid stimulus trains, we used the kinetically rapid opsin Chronos (Klapoetke, Murata et al. 2014). Example traces are shown in Fig. 8B. The relatively robust paired pulse facilitation in WT mice for both dSPN and iSPN (Fig. 8C,D; black) is substantially reduced by DA depletion (Fig. 8C,D; red). This effect is most significant at 10 Hz and 20 Hz for dSPN and iSPN. The 33 Hz stimuli had the same trend but the effect was not significant, though higher rates are more limiting, even for kinetically rapid opsins (Klapoetke, Murata et al. 2014). Surprisingly, DA depletion in cTED mice did not show such an effect, but rather paired pulse facilitation that is observed at baseline was maintained in these mice (Fig. 8C,D; blue and magenta). Thus, the presynaptic thalamic paired pulse facilitation at PF to SPN synapses is maintained in cTED mice despite the loss of DA. We thus hypothesize that maintenance of normal thalamostriatal transmission during trains might play a role in maintaining normal motor system operation and preserving paw preference even after loss of DA neurons in this model.

## DISCUSSION

In this study, we report on mice with normal paw reaching behavior despite a unilateral depletion of dorsal striatal DA. Specifically, we show that the normal paw reaching behavior is induced by transiently increased striatal DA release at the onset of the waking cycle (either via transgenic cTED or via eDREADD activation in the iTED model). We also determined that the transient increase in DA is required prior to injection of 6-OHDA, indicating a priming step for induction of the plasticity, as well as for seven days following the injection. Behavioral preservation was accompanied by a morphological preservation of mature spine density and dendritic arborization specifically in dSPNs, and not iSPNs, of both cTED and iTED mice. Both the behavioral and morphological preservation were lost with pharmacological blockade of the TED enhanced striatal D1 receptor activation (D1 antagoinist administered directly to the striatum prior to and during DA depletion). The tight correlation between preserved dSPN morphology and normal motor behavior after DA depletion is consistent with preserved striatal output (i.e. direct over indirect pathway) (Stephens, Mueller et al. 2005, Smith, Raju et al. 2009). However, a detailed examination of M1 corticostriatal input showed a profound loss of both IT-type and PT-type corticostriatal input to SPNs in both WT and cTED mice. Parafasicular thalamostriatal input strength were preserved in both WT and cTED mice, while short-term plasticity was preserved specifically in only the cTED model. Together, these findings suggest that preserved motor behavior is associated with the maintenance of dSPN morphology and the preservation of thalamostriatal synaptic function, including paired pulse facilitation. Future experiments will be required to determine whether these circuit adaptations in DA-depleted TED mice contribute causally to the preservation of motor function.

### Alterations in Excitatory Input

M1 provides a major source of excitatory input to the dorsolateral striatum (Ding, Peterson et al. 2008) and therefore was a likely source of input to the preserved mature spines on dSPNs of cTED and iTED mice after DA depletion. Consistent with previous reports (Fieblinger, Graves et al. 2014), 6-OHDA treated controls show reduced mature dSPN dendritic spine density. In contrast, mature spine density was preserved in dSPNs of DA-depleted TED mice (both cTED and iTED), potentially resulting from the maturation of immature dSPN spines which are elevated in non-depleted cTED and iTED mice during times when striatal DA release is elevated (Fig. S3A,B). DA loss also increased dSPN excitability in WT mice, as reported previously (Fieblinger, Graves et al. 2014, Maurice, Liberge et al. 2015, Ketzef, Spigolon et al. 2017). Although reduced glutamatergic input from M1 may contribute to spine loss in WT mice, increased excitability of dSPNs may represent the homeostatic response to diminished excitatory input in WT mice consistent with published reports. However, dSPNs of depleted cTED mice exhibited an excitability more similar to non-depleted WT and cTED mice. Lost input could have triggered a homeostatic drive resulting in excitability changes, though observations when taken together, suggest that in cTED mice, the reluctance to change excitability may be a result of excitatory drive strengthened from sources other than M1, such as other cortical areas or thalamus. While we hypothesize that transient elevation of DA release prevents these deleterious physiological adaptations, our work also suggests that further investigation into the source of the maintained mature spines on dSPNs is warranted.

In iSPNs of TED mice, mature spine density and excitability were unchanged relative to WT and control mice. These findings, consistent with previous reports (Suarez, Solis et al. 2020), suggest that elevated DA does not elicit a homeostatic response in iSPNs, as opposed to dSPNs. Because iSPNs were identified as D1R-tdTomato negative neurons, a small number of non-iSPN, including the cholinergic and parvalbumin+ interneurons, may have been included despite exclusion criteria based on capacitance, intrinsic current kinetics, and firing properties.

The induction of plasticity that manifests as preserved motor behavior in TED mice depends on the extent of dopamine depletion as mfb lesions which deplete DA from both the nigrostriatal and VTA-nucleus accumbens circuits did not result in preserved motor behavior (data not shown). Further, D1R is implicated in the induction of long-term potentiation (LTP) and D2R is associated with the induction of long-term depression (LTD) (Kerr and Wickens 2001, Lovinger 2010, Paille, Picconi et al. 2010). Also, the induction and maintenance of LTP and LTD require different levels of DA (Paille, Picconi et al. 2010). A complete loss of striatal DA leads to a complete loss of LTP and LTD, whereas a reduction in DA levels limited to the dorsal striatum has more selective effects. It was shown that alterations in the measured intrinsic membrane properties (excitability) of striatal spiny neurons depend on the induction of LTP and LTD (Paille, Picconi et al. 2010). It is quite possible that intrastriatal depletion exhibits changes linked with LTP and hence excitability changes in dSPNs, and that other models such as mfb lesions are more likely to cause excitability changes in iSPNs.

Our connectivity data show that in control cohorts, IT-type projections provide the strongest synaptic input to dSPNs and iSPNs, roughly three times stronger than synaptic input from PT-type neurons. Consistent with prior experiments, IT-type inputs from M1 slightly favor iSPNs over dSPNs (Wall, De La Parra et al. 2013). After DA depletion, cortical inputs measured in control cohorts were substantially lost (>70%) with the reduction notably more severe for PT-type inputs than IT-type inputs. This differs from more modest alterations in cortical input in other DA depletion models (Parker, Lalive et al. 2016). cTED mice also showed significant reductions in IT- and PT-type inputs to dSPNs as well as iSPNs, inconsistent with the expectation that normal corticostriatal input would be necessary for the observed behavioral preservation in the paw reaching behavior and normal mature spine density in dSPNs. Indeed, despite the loss of overall input strength from M1, the dSPN/iSPN input ratio is not disrupted following depletion (Fig. 6E,I). Because these experiments were restricted to M1-derived corticostriatal projections, adaptive changes in dSPNs of TED mice resulting from enhanced input of other cortical regions, including M2 and somatosensory cortex that converge on the same dSPNs (Hooks, Papale et al. 2018), cannot be excluded. In contrast, thalamostriatal inputs are preserved in both control and cTED mice, consistent with previous studies where DA depletion reduced input signals from the PF nucleus to the dSPNs (Parker, Lalive et al. 2016, Tanimura, Shen et al. 2022).

The preservation of PF thalamostriatal input strength in both control and TED mice cannot alone explain the selective sparing of motor output in the TED model. The paired pulse experiments revealed a reduced paired-pulse ratio (PPR) in the DA-depleted WT condition (Fig. 8C-D), suggesting that there is an increased presynaptic probability of glutamate release at the PF-SPN synapse (for both direct and indirect pathways). But because the overall thalamostriatal input strength is not changed for DA-depleted WT mice (Fig. 7), this suggests some inputs are lost, while remaining ones have enhanced release probability. In contrast, we observed that synaptic facilitation, as occurs during thalamic bursting (McCormick and Pape 1990, Sherman and Guillery 2006), is preserved in depleted cTED mice at the same level as naïve WT and cTED mice. This suggests that the probability of glutamate release is maintained in a like-normal condition, possibly due to prevention of thalamostriatal synapse remodeling. Thus, the preservation of thalamostriatal transmission during bursts of activity may contribute to maintenance of normal motor behavior in cTED mice following DA depletion.

Glutamatergic excitatory projections terminate at the spine heads, where glutamatergic receptors, including N-methyl-D-aspartate (NMDA), are localized (Kemp and Powell 1971, Bolam, Hanley et al. 2000, Vastagh, Gardoni et al. 2012). Cortical input targets spines, while thalamic input contacts both spines and dendritic shafts (Smith, Raju et al. 2009). Our results reveal an increase in NMDAR mediated currents in DA depleted conditions (Fig. 7J). These results are consistent with several other studies, where following loss of 75% of TH+ axons in the striatum resulted in an increase in the NMDA-NR2A subunit in the postsynaptic component (Chase, Oh et al. 1998, Paille, Picconi et al. 2010). D1 receptor activation leads to phosphorylation-dependent trafficking of NMDAR to the dendritic membrane (Hallett, Dunah et al. 2005). The increased NMDAR currents observed after DA depletion, as shown in these studies (Fig. 7J), may lead to greater postsynaptic calcium (Jones-Tabah, Mohammad et al. 2021). This would activate downstream second messenger pathways to increase synaptic strength, consistent with published reports (Gerfen, Miyachi et al. 2002), including studies with FRET-based biosensors (Jones-Tabah, Mohammad et al. 2020). In cTED mice, the NMDAR/AMPAR ratio is lower than control and significantly lower than depleted cTED mice. Thus, in the depleted cTED mice, the plasticity produces a decrease in the NMDAR mediated currents and a possible restoration of normal function. Overall, based on a current understanding in the field, the balance between dSPN and iSPN activity regulates voluntary movement through two opposing pathways in the brain’s basal ganglia: the direct (“go”) pathway and the indirect (“stop”) pathway (Kravitz, Freeze et al. 2010, Cui, Jun et al. 2013, Panigrahi, Martin et al. 2015, Markowitz, Gillis et al. 2018, Parker, Marshall et al. 2018). The direct pathway, via dSPNs, promotes and initiates movement. In Parkinson’s disease, without enough DA, dSPN-driven “go” signals are reduced and iSPN-driven “stop” signals become hyperactive, creating an imbalance in the movement regulation. The preservation of paw-reaching behavior suggests maintenance of functional output from the basal ganglia circuits despite substantial DA depletion. Putative mechanisms for maintaining normal motor output include (a) preservation and strengthening of other inputs to the dSPN spines, such as other cortical areas (including S1 and M2) or other motor thalamic regions (such as CL), or (b) preservation of thalamic input during trains of stimuli. Because of the transgenic tools for circuit analysis available in mouse, this should be a fruitful system for identifying circuit loci where movement deficits are induced and could be rescued (McGregor and Nelson 2019).

### Comparison of Mouse Models of DA Loss

Here, we use a dorsal striatal lesion rodent model of PD (Ungerstedt 1968, Brundin, Isacson et al. 1986, Przedborski, Levivier et al. 1995, Lee, Sauer et al. 1996, Kirik, Rosenblad et al. 1998, Barneoud, Descombris et al. 2000, Iancu, Mohapel et al. 2005, Tieu 2011) that limits DA depletion to the nigrostriatal pathway, thus better modeling the anatomical location of DA loss in human PD during the earlier stages (Boix, Padel et al. 2015). This model provides a more progressive and stable depletion of DA, replicating motor and non-motor deficits consistent with the human disease (Tieu 2011, Mendes-Pinheiro, Soares-Cunha et al. 2021). Other commonly used 6-OHDA models target the mfb (Carman, Gage et al. 1991, Dentresangle, Le Cavorsin et al. 2001, Day, Wang et al. 2006, Paille, Picconi et al. 2010, Parker, Lalive et al. 2016, Mastro, Zitelli et al. 2017), which rapidly lesions the entire midbrain DA system, including loss of the VTA to nucleus accumbens, or direct injection into the SNpC which is challenged by the close proximity of the VTA (Deumens, Blokland et al. 2002). The effect of mfb injections is similar to intrastriatal injections at similar doses (Lundblad, Picconi et al. 2004). mfb lesions produce extensive degeneration that may not accurately model the earlier stages of human PD (Boix, Padel et al. 2015). Confirmation of our findings in rodent or monkey PD models (ideally with means to elevate dopamine neuron activity prior to and during degeneration) that produce only nigrostriatal degeneration, have neuropathology like alpha synuclein inclusions, or mimic a slower degeneration as is seen in a subset of PD patients, will provide additional insight into the clinical utility of these findings.

### Implications for Early Treatment

Our findings provide a new strategy to improve therapeutic outcomes for motor symptoms of PD. Early detection either through genetic screening or detection of prodromal symptoms coupled with a TED-like treatment to transiently elevate nigrostriatal dopamine neuron activity could slow the onset of motor symptoms. Interestingly, exercise, which is employed as an adjunct treatment for PD because of its neuroprotective and restorative effects (Palasz, Niewiadomski et al. 2019), elevates striatal DA and promotes spine enhancement within the mammalian brain (Hou, Chen et al. 2017), including on dSPNs and iSPNs in the MPTP mouse model of PD (Toy, Petzinger et al. 2014)). The spine enhancement may be mediated through neurotrophins, which are secreted from both cortico- and nigrostriatal afferents (Conner, Lauterborn et al. 1997, Baquet, Gorski et al. 2004, Baydyuk and Xu 2014). Neurotrophins play a critical role in neuronal growth, maintenance, and plasticity (Chao 2003, Park and Poo 2013), and have been shown to be neuroprotective in PD animal models (Nagahara and Tuszynski 2011). Thus, mechanisms we identified here may be similarly engaged by the exercises that benefit PD patients. Additional therapeutic strategies that could potentially induce this striatal plasticity mechanism at early stages of PD include identifying and manipulating VGLUT3^+^ neurons that underlie the hyperdopaminergic phenotype in the KO. Lastly, findings presented here may have implications beyond motor function, for example, midbrain DA loss underlying mood, sensory and cognitive symptoms commonly co-morbid in PD (Marras and Chaudhuri 2016, Goldman, Vernaleo et al. 2018).

## ACKNOWLEDGMENTS

We thank Dr. Myung-chul Noh for help with electrophysiological recordings and comments on the manuscript; Drs. David Ferreira and Cynthia Arokiaraj for experimental design input and comments on the manuscript; Dr. Aryn Gittis for technical advice; Yan Dong for technical advice and comments on the manuscript; Cole Boillat, Daniel Butz, Jenna Nemschoff, Zane Hamden, Amy Liu, Maximillion Junge, Tulika Malik, Nikhill Khongbantabam, Thalia, Petroulakis, Randi Wilk, and Gopika Pillai for counting spines and scoring behavior; Jeremy Gedeon for technical assistance; Sean Paul Williams and Kelly Corrigan for mouse colony maintenance; Suh Jin Lee, Kelly Corrigan, Rui Peixoto, Zachary Freyberg, and Jennifer Magnusson for comments on the manuscript. Center for Biological Imaging and Simon Watkins for confocal microscope support; Suh Jin Lee for illustration in Figure S1D. Funding was T32NS086749 (J.C.B.), R01NS082650 (R.P.S.), DoD-Army: W81XWH-171-0386 (R.P.S.), a UPBI NeuroDiscovery Pilot Grant (R.P.S.), R01NS103993 (B.M.H.), and a Whitehall Foundation award (B.M.H.).

## Author Contributions

R.P.S. conceived the study; R.P.S., J.C.B., G. P. S., and B.M.H designed experiments; J.C.B., G.P.S., Z.H. and D.A.H. performed the experiments, J.C.B., G.P.S., B.M.H, Z.H., D.J.H., D.A.H and R.P.S analyzed and interpreted the data; J.C.B., G.P.S., B.M.H. and R.P.S. wrote the paper.

## REFERENCES

Ade, K. K., Y. Wan, M. Chen, B. Gloss and N. Calakos (2011). “An Improved BAC Transgenic Fluorescent Reporter Line for Sensitive and Specific Identification of Striatonigral Medium Spiny Neurons.” Front Syst Neurosci 5: 32.

Albin, R. L., A. B. Young and J. B. Penney (1989). “The functional anatomy of basal ganglia disorders.” Trends Neurosci 12(10): 366–375.

Alexander, G. E., M. R. DeLong and P. L. Strick (1986). “Parallel organization of functionally segregated circuits linking basal ganglia and cortex.” Annu Rev Neurosci 9: 357–381.

Armbruster, B. N., X. Li, M. H. Pausch, S. Herlitze and B. L. Roth (2007). “Evolving the lock to fit the key to create a family of G protein-coupled receptors potently activated by an inert ligand.” Proc Natl Acad Sci U S A 104(12): 5163–5168.

Baquet, Z. C., J. A. Gorski and K. R. Jones (2004). “Early striatal dendrite deficits followed by neuron loss with advanced age in the absence of anterograde cortical brain-derived neurotrophic factor.” J Neurosci 24(17): 4250–4258.

Barneoud, P., E. Descombris, N. Aubin and D. N. Abrous (2000). “Evaluation of simple and complex sensorimotor behaviours in rats with a partial lesion of the dopaminergic nigrostriatal system.” Eur J Neurosci 12(1): 322–336.

Baydyuk, M. and B. Xu (2014). “BDNF signaling and survival of striatal neurons.” Front Cell Neurosci 8: 254.

Boix, J., T. Padel and G. Paul (2015). “A partial lesion model of Parkinson’s disease in mice--characterization of a 6-OHDA-induced medial forebrain bundle lesion.” Behav Brain Res 284: 196–206.

Bolam, J. P., J. J. Hanley, P. A. Booth and M. D. Bevan (2000). “Synaptic organisation of the basal ganglia.” J Anat 196 (Pt 4)(Pt 4): 527–542.

Brundin, P., O. Isacson, F. H. Gage and A. Bjorklund (1986). “Intrastriatal grafting of dopamine-containing neuronal cell suspensions: effects of mixing with target or non-target cells.” Brain Res 389(1-2): 77–84.

Carman, L. S., F. H. Gage and C. W. Shults (1991). “Partial lesion of the substantia nigra: relation between extent of lesion and rotational behavior.” Brain Res 553(2): 275–283.

Chao, M. V. (2003). “Neurotrophins and their receptors: a convergence point for many signalling pathways.” Nat Rev Neurosci 4(4): 299–309.

Chase, T. N., J. D. Oh and P. J. Blanchet (1998). “Neostriatal mechanisms in Parkinson’s disease.” Neurology 51(2 Suppl 2): S30–35.

Conner, J. M., J. C. Lauterborn, Q. Yan, C. M. Gall and S. Varon (1997). “Distribution of brain-derived neurotrophic factor (BDNF) protein and mRNA in the normal adult rat CNS: evidence for anterograde axonal transport.” J Neurosci 17(7): 2295–2313.

Cui, G., S. B. Jun, X. Jin, M. D. Pham, S. S. Vogel, D. M. Lovinger and R. M. Costa (2013). “Concurrent activation of striatal direct and indirect pathways during action initiation.” Nature 494(7436): 238–242.

Currie, S. P., J. J. Ammer, B. Premchand, J. Dacre, Y. Wu, C. Eleftheriou, M. Colligan, T. Clarke, L. Mitchell, A. A. Faisal, M. H. Hennig and I. Duguid (2022). “Movement-specific signaling is differentially distributed across motor cortex layer 5 projection neuron classes.” Cell Rep 39(6): 110801.

Day, M., Z. Wang, J. Ding, X. An, C. A. Ingham, A. F. Shering, D. Wokosin, E. Ilijic, Z. Sun, A. R. Sampson, E. Mugnaini, A. Y. Deutch, S. R. Sesack, G. W. Arbuthnott and D. J. Surmeier (2006). “Selective elimination of glutamatergic synapses on striatopallidal neurons in Parkinson disease models.” Nat Neurosci 9(2): 251–259.

Dentresangle, C., M. Le Cavorsin, M. Savasta and V. Leviel (2001). “Increased extracellular DA and normal evoked DA release in the rat striatum after a partial lesion of the substantia nigra.” Brain Res 893(1-2): 178–185.

Deumens, R., A. Blokland and J. Prickaerts (2002). “Modeling Parkinson’s disease in rats: an evaluation of 6-OHDA lesions of the nigrostriatal pathway.” Exp Neurol 175(2): 303–317.

Ding, J., J. D. Peterson and D. J. Surmeier (2008). “Corticostriatal and thalamostriatal synapses have distinctive properties.” J Neurosci 28(25): 6483–6492.

Divito, C. B., K. Steece-Collier, D. T. Case, S. P. Williams, J. A. Stancati, L. Zhi, M. E. Rubio, C. E. Sortwell, T. J. Collier, D. Sulzer, R. H. Edwards, H. Zhang and R. P. Seal (2015). “Loss of VGLUT3 Produces Circadian-Dependent Hyperdopaminergia and Ameliorates Motor Dysfunction and l-Dopa-Mediated Dyskinesias in a Model of Parkinson’s Disease.” J Neurosci 35(45): 14983–14999.

Fieblinger, T., S. M. Graves, L. E. Sebel, C. Alcacer, J. L. Plotkin, T. S. Gertler, C. S. Chan, M. Heiman, P. Greengard, M. A. Cenci and D. J. Surmeier (2014). “Cell type-specific plasticity of striatal projection neurons in parkinsonism and L-DOPA-induced dyskinesia.” Nat Commun 5: 5316.

Gagnon, D., S. Petryszyn, M. G. Sanchez, C. Bories, J. M. Beaulieu, Y. De Koninck, A. Parent and M. Parent (2017). “Striatal Neurons Expressing D1 and D2 Receptors are Morphologically Distinct and Differently Affected by Dopamine Denervation in Mice.” Sci Rep 7: 41432.

Gerfen, C. R., T. M. Engber, L. C. Mahan, Z. Susel, T. N. Chase, F. J. Monsma, Jr. and D. R. Sibley (1990). “D1 and D2 dopamine receptor-regulated gene expression of striatonigral and striatopallidal neurons.” Science 250(4986): 1429–1432.

Gerfen, C. R., S. Miyachi, R. Paletzki and P. Brown (2002). “D1 dopamine receptor supersensitivity in the dopamine-depleted striatum results from a switch in the regulation of ERK1/2/MAP kinase.” J Neurosci 22(12): 5042–5054.

Gerfen, C. R., R. Paletzki and N. Heintz (2013). “GENSAT BAC cre-recombinase driver lines to study the functional organization of cerebral cortical and basal ganglia circuits.” Neuron 80(6): 1368–1383.

Goldman, J. G., B. A. Vernaleo, R. Camicioli, N. Dahodwala, R. D. Dobkin, T. Ellis, J. E. Galvin, C. Marras, J. Edwards, J. Fields, R. Golden, J. Karlawish, B. Levin, L. Shulman, G. Smith, C. Tangney, C. A. Thomas, A. I. Troster, E. Y. Uc, N. Coyan, C. Ellman, M. Ellman, C. Hoffman, S. Hoffman and D. Simmonds (2018). “Cognitive impairment in Parkinson’s disease: a report from a multidisciplinary symposium on unmet needs and future directions to maintain cognitive health.” NPJ Parkinsons Dis 4: 19.

Gras, C., B. Amilhon, E. M. Lepicard, O. Poirel, J. Vinatier, M. Herbin, S. Dumas, E. T. Tzavara, M. R. Wade, G. G. Nomikos, N. Hanoun, F. Saurini, M. L. Kemel, B. Gasnier, B. Giros and S. El Mestikawy (2008). “The vesicular glutamate transporter VGLUT3 synergizes striatal acetylcholine tone.” Nat Neurosci 11(3): 292–300.

Hallett, P. J., A. W. Dunah, P. Ravenscroft, S. Zhou, E. Bezard, A. R. Crossman, J. M. Brotchie and D. G. Standaert (2005). “Alterations of striatal NMDA receptor subunits associated with the development of dyskinesia in the MPTP-lesioned primate model of Parkinson’s disease.” Neuropharmacology 48(4): 503–516.

Harris, K. D. and G. M. Shepherd (2015). “The neocortical circuit: themes and variations.” Nat Neurosci 18(2): 170–181.

Hintiryan, H., N. N. Foster, I. Bowman, M. Bay, M. Y. Song, L. Gou, S. Yamashita, M. S. Bienkowski, B. Zingg, M. Zhu, X. W. Yang, J. C. Shih, A. W. Toga and H. W. Dong (2016). “The mouse cortico-striatal projectome.” Nat Neurosci 19(8): 1100–1114.

Hooks, B. M., J. Y. Lin, C. Guo and K. Svoboda (2015). “Dual-Channel Circuit Mapping Reveals Sensorimotor Convergence in the Primary Motor Cortex.” J Neurosci 35(10): 4418–4426.

Hooks, B. M., A. E. Papale, R. F. Paletzki, M. W. Feroze, B. S. Eastwood, J. J. Couey, J. Winnubst, J. Chandrashekar and C. R. Gerfen (2018). “Topographic precision in sensory and motor corticostriatal projections varies across cell type and cortical area.” Nat Commun 9(1): 3549.

Hou, L., W. Chen, X. Liu, D. Qiao and F. M. Zhou (2017). “Exercise-Induced Neuroprotection of the Nigrostriatal Dopamine System in Parkinson’s Disease.” Front Aging Neurosci 9: 358.

Huang, Y. H., Y. Lin, P. Mu, B. R. Lee, T. E. Brown, G. Wayman, H. Marie, W. Liu, Z. Yan, B. A. Sorg, O. M. Schluter, R. S. Zukin and Y. Dong (2009). “In vivo cocaine experience generates silent synapses.” Neuron 63(1): 40–47.

Iancu, R., P. Mohapel, P. Brundin and G. Paul (2005). “Behavioral characterization of a unilateral 6-OHDA-lesion model of Parkinson’s disease in mice.” Behav Brain Res 162(1): 1–10.

Jones-Tabah, J., H. Mohammad, S. Hadj-Youssef, L. E. H. Kim, R. D. Martin, F. Benaliouad, J. C. Tanny, P. B. S. Clarke and T. E. Hebert (2020). “Dopamine D1 receptor signalling in dyskinetic Parkinsonian rats revealed by fiber photometry using FRET-based biosensors.” Sci Rep 10(1): 14426.

Jones-Tabah, J., H. Mohammad, E. G. Paulus, P. B. S. Clarke and T. E. Hebert (2021). “The Signaling and Pharmacology of the Dopamine D1 Receptor.” Front Cell Neurosci 15: 806618.

Kemp, J. M. and T. P. Powell (1971). “The termination of fibres from the cerebral cortex and thalamus upon dendritic spines in the caudate nucleus: a study with the Golgi method.” Philos Trans R Soc Lond B Biol Sci 262(845): 429–439.

Kerr, J. N. and J. R. Wickens (2001). “Dopamine D-1/D-5 receptor activation is required for long-term potentiation in the rat neostriatum in vitro.” J Neurophysiol 85(1): 117–124.

Ketzef, M., G. Spigolon, Y. Johansson, A. Bonito-Oliva, G. Fisone and G. Silberberg (2017). “Dopamine Depletion Impairs Bilateral Sensory Processing in the Striatum in a Pathway-Dependent Manner.” Neuron 94(4): 855–865 e855.

Kirik, D., C. Rosenblad and A. Bjorklund (1998). “Characterization of behavioral and neurodegenerative changes following partial lesions of the nigrostriatal dopamine system induced by intrastriatal 6-hydroxydopamine in the rat.” Exp Neurol 152(2): 259–277.

Klapoetke, N. C., Y. Murata, S. S. Kim, S. R. Pulver, A. Birdsey-Benson, Y. K. Cho, T. K. Morimoto, A. S. Chuong, E. J. Carpenter, Z. Tian, J. Wang, Y. Xie, Z. Yan, Y. Zhang, B. Y. Chow, B. Surek, M. Melkonian, V. Jayaraman, M. Constantine-Paton, G. K. Wong and E. S. Boyden (2014). “Independent optical excitation of distinct neural populations.” Nat Methods 11(3): 338–346.

Krashes, M. J., S. Koda, C. Ye, S. C. Rogan, A. C. Adams, D. S. Cusher, E. Maratos-Flier, B. L. Roth and B. B. Lowell (2011). “Rapid, reversible activation of AgRP neurons drives feeding behavior in mice.” J Clin Invest 121(4): 1424–1428.

Kravitz, A. V., B. S. Freeze, P. R. Parker, K. Kay, M. T. Thwin, K. Deisseroth and A. C. Kreitzer (2010). “Regulation of parkinsonian motor behaviours by optogenetic control of basal ganglia circuitry.” Nature 466(7306): 622–626.

Lanciego, J. L., N. Luquin and J. A. Obeso (2012). “Functional neuroanatomy of the basal ganglia.” Cold Spring Harb Perspect Med 2(12): a009621.

Lee, B. R., Y. Y. Ma, Y. H. Huang, X. Wang, M. Otaka, M. Ishikawa, P. A. Neumann, N. M. Graziane, T. E. Brown, A. Suska, C. Guo, M. K. Lobo, S. R. Sesack, M. E. Wolf, E. J. Nestler, Y. Shaham, O. M. Schluter and Y. Dong (2013). “Maturation of silent synapses in amygdala-accumbens projection contributes to incubation of cocaine craving.” Nat Neurosci 16(11): 1644–1651.

Lee, C. S., H. Sauer and A. Bjorklund (1996). “Dopaminergic neuronal degeneration and motor impairments following axon terminal lesion by instrastriatal 6-hydroxydopamine in the rat.” Neuroscience 72(3): 641–653.

Lee, J., W. Wang and B. L. Sabatini (2020). “Anatomically segregated basal ganglia pathways allow parallel behavioral modulation.” Nat Neurosci 23(11): 1388–1398.

Lin, J. Y., P. M. Knutsen, A. Muller, D. Kleinfeld and R. Y. Tsien (2013). “ReaChR: a red-shifted variant of channelrhodopsin enables deep transcranial optogenetic excitation.” Nat Neurosci 16(10): 1499–1508.

Lisman, J. E., J. M. Fellous and X. J. Wang (1998). “A role for NMDA-receptor channels in working memory.” Nat Neurosci 1(4): 273–275.

Lovinger, D. M. (2010). “Neurotransmitter roles in synaptic modulation, plasticity and learning in the dorsal striatum.” Neuropharmacology 58(7): 951–961.

Lundblad, M., B. Picconi, H. Lindgren and M. A. Cenci (2004). “A model of L-DOPA-induced dyskinesia in 6-hydroxydopamine lesioned mice: relation to motor and cellular parameters of nigrostriatal function.” Neurobiol Dis 16(1): 110–123.

Markowitz, J. E., W. F. Gillis, C. C. Beron, S. Q. Neufeld, K. Robertson, N. D. Bhagat, R. E. Peterson, E. Peterson, M. Hyun, S. W. Linderman, B. L. Sabatini and S. R. Datta (2018). “The Striatum Organizes 3D Behavior via Moment-to-Moment Action Selection.” Cell 174(1): 44–58 e17.

Marras, C. and K. R. Chaudhuri (2016). “Nonmotor features of Parkinson’s disease subtypes.” Mov Disord 31(8): 1095–1102.

Mastro, K. J., K. T. Zitelli, A. M. Willard, K. H. Leblanc, A. V. Kravitz and A. H. Gittis (2017). “Cell-specific pallidal intervention induces long-lasting motor recovery in dopamine-depleted mice.” Nat Neurosci 20(6): 815–823.

Maurice, N., M. Liberge, F. Jaouen, S. Ztaou, M. Hanini, J. Camon, K. Deisseroth, M. Amalric, L. Kerkerian-Le Goff and C. Beurrier (2015). “Striatal Cholinergic Interneurons Control Motor Behavior and Basal Ganglia Function in Experimental Parkinsonism.” Cell Rep 13(4): 657–666.

McCormick, D. A. and H. C. Pape (1990). “Properties of a hyperpolarization-activated cation current and its role in rhythmic oscillation in thalamic relay neurones.” Journal of Physiology 431: 291–318.

McGregor, M. M. and A. B. Nelson (2019). “Circuit Mechanisms of Parkinson’s Disease.” Neuron 101(6): 1042–1056.

Mendes-Pinheiro, B., C. Soares-Cunha, A. Marote, E. Loureiro-Campos, J. Campos, S. Barata-Antunes, D. Monteiro-Fernandes, D. Santos, S. Duarte-Silva, L. Pinto and A. Jose Salgado (2021). “Unilateral Intrastriatal 6-Hydroxydopamine Lesion in Mice: A Closer Look into Non-Motor Phenotype and Glial Response.” Int J Mol Sci 22(21).

Nagahara, A. H. and M. H. Tuszynski (2011). “Potential therapeutic uses of BDNF in neurological and psychiatric disorders.” Nat Rev Drug Discov 10(3): 209–219.

Paille, V., B. Picconi, V. Bagetta, V. Ghiglieri, C. Sgobio, M. Di Filippo, M. T. Viscomi, C. Giampa, F. R. Fusco, F. Gardoni, G. Bernardi, P. Greengard, M. Di Luca and P. Calabresi (2010). “Distinct levels of dopamine denervation differentially alter striatal synaptic plasticity and NMDA receptor subunit composition.” J Neurosci 30(42): 14182–14193.

Palasz, E., W. Niewiadomski, A. Gasiorowska, A. Wysocka, A. Stepniewska and G. Niewiadomska (2019). “Exercise-Induced Neuroprotection and Recovery of Motor Function in Animal Models of Parkinson’s Disease.” Front Neurol 10: 1143.

Panigrahi, B., K. A. Martin, Y. Li, A. R. Graves, A. Vollmer, L. Olson, B. D. Mensh, A. Y. Karpova and J. T. Dudman (2015). “Dopamine Is Required for the Neural Representation and Control of Movement Vigor.” Cell 162(6): 1418–1430.

Park, H. and M. M. Poo (2013). “Neurotrophin regulation of neural circuit development and function.” Nat Rev Neurosci 14(1): 7–23.

Park, J., J. W. Phillips, J. Z. Guo, K. A. Martin, A. W. Hantman and J. T. Dudman (2022). “Motor cortical output for skilled forelimb movement is selectively distributed across projection neuron classes.” Sci Adv 8(10): eabj5167.

Parker, J. G., J. D. Marshall, B. Ahanonu, Y. W. Wu, T. H. Kim, B. F. Grewe, Y. Zhang, J. Z. Li, J. B. Ding, M. D. Ehlers and M. J. Schnitzer (2018). “Diametric neural ensemble dynamics in parkinsonian and dyskinetic states.” Nature 557(7704): 177–182.

Parker, P. R., A. L. Lalive and A. C. Kreitzer (2016). “Pathway-Specific Remodeling of Thalamostriatal Synapses in Parkinsonian Mice.” Neuron 89(4): 734–740.

Pasquereau, B., M. R. DeLong and R. S. Turner (2016). “Primary motor cortex of the parkinsonian monkey: altered encoding of active movement.” Brain 139(Pt 1): 127–143.

Petreanu, L., D. Huber, A. Sobczyk and K. Svoboda (2007). “Channelrhodopsin-2-assisted circuit mapping of long-range callosal projections.” Nat Neurosci 10(5): 663–668.

Petreanu, L., T. Mao, S. M. Sternson and K. Svoboda (2009). “The subcellular organization of neocortical excitatory connections.” Nature 457(7233): 1142–1145.

Phillips, J. W., A. Schulmann, E. Hara, J. Winnubst, C. Liu, V. Valakh, L. Wang, B. C. Shields, W. Korff, J. Chandrashekar, A. L. Lemire, B. Mensh, J. T. Dudman, S. B. Nelson and A. W. Hantman (2019). “A repeated molecular architecture across thalamic pathways.” Nat Neurosci 22(11): 1925–1935.

Popescu, A. T., A. A. Saghyan and D. Pare (2007). “NMDA-dependent facilitation of corticostriatal plasticity by the amygdala.” Proc Natl Acad Sci U S A 104(1): 341–346.

Prescott, S. A. and Y. De Koninck (2002). “Four cell types with distinctive membrane properties and morphologies in lamina I of the spinal dorsal horn of the adult rat.” J Physiol 539(Pt 3): 817–836.

Przedborski, S., M. Levivier, H. Jiang, M. Ferreira, V. Jackson-Lewis, D. Donaldson and D. M. Togasaki (1995). “Dose-dependent lesions of the dopaminergic nigrostriatal pathway induced by intrastriatal injection of 6-hydroxydopamine.” Neuroscience 67(3): 631–647.

Seal, R. P., O. Akil, E. Yi, C. M. Weber, L. Grant, J. Yoo, A. Clause, K. Kandler, J. L. Noebels, E. Glowatzki, L. R. Lustig and R. H. Edwards (2008). “Sensorineural deafness and seizures in mice lacking vesicular glutamate transporter 3.” Neuron 57(2): 263–275.

Shepherd, G. M. (2013). “Corticostriatal connectivity and its role in disease.” Nat Rev Neurosci 14(4): 278–291.

Shepherd, G. M., T. A. Pologruto and K. Svoboda (2003). “Circuit analysis of experience-dependent plasticity in the developing rat barrel cortex.” Neuron 38(2): 277–289.

Sherman, S. M. and R. W. Guillery (2006). Exploring the thalamus and its role in cortical function. Cambridge, Mass., MIT Press.

Shinotsuka, T., Y. R. Tanaka, S. I. Terada, N. Hatano and M. Matsuzaki (2023). “Layer 5 Intratelencephalic Neurons in the Motor Cortex Stably Encode Skilled Movement.” J Neurosci 43(43): 7130–7148.

Smeal, R. M., K. A. Keefe and K. S. Wilcox (2008). “Differences in excitatory transmission between thalamic and cortical afferents to single spiny efferent neurons of rat dorsal striatum.” Eur J Neurosci 28(10): 2041–2052.

Smith, Y., M. D. Bevan, E. Shink and J. P. Bolam (1998). “Microcircuitry of the direct and indirect pathways of the basal ganglia.” Neuroscience 86(2): 353–387.

Smith, Y., D. Raju, B. Nanda, J. F. Pare, A. Galvan and T. Wichmann (2009). “The thalamostriatal systems: anatomical and functional organization in normal and parkinsonian states.” Brain Res Bull 78(2-3): 60–68.

Smith, Y. and R. Villalba (2008). “Striatal and extrastriatal dopamine in the basal ganglia: an overview of its anatomical organization in normal and Parkinsonian brains.” Mov Disord 23 Suppl 3: S534–547.

Smith, Y., R. M. Villalba and D. V. Raju (2009). “Striatal spine plasticity in Parkinson’s disease: pathological or not?” Parkinsonism Relat Disord 15 Suppl 3(Suppl 3): S156–161.

Stephens, B., A. J. Mueller, A. F. Shering, S. H. Hood, P. Taggart, G. W. Arbuthnott, J. E. Bell, L. Kilford, A. E. Kingsbury, S. E. Daniel and C. A. Ingham (2005). “Evidence of a breakdown of corticostriatal connections in Parkinson’s disease.” Neuroscience 132(3): 741–754.

Suarez, L. M., O. Solis, A. Sanz-Magro, S. Alberquilla and R. Moratalla (2020). “Dopamine D1 Receptors Regulate Spines in Striatal Direct-Pathway and Indirect-Pathway Neurons.” Mov Disord 35(10): 1810–1821.

Tanimura, A., W. Shen, D. Wokosin and D. J. Surmeier (2022). “Pathway-Specific Remodeling of Thalamostriatal Synapses in a Mouse Model of Parkinson’s Disease.” Mov Disord 37(6): 1164–1174.

Tieu, K. (2011). “A guide to neurotoxic animal models of Parkinson’s disease.” Cold Spring Harb Perspect Med 1(1): a009316.

Toy, W. A., G. M. Petzinger, B. J. Leyshon, G. K. Akopian, J. P. Walsh, M. V. Hoffman, M. G. Vuckovic and M. W. Jakowec (2014). “Treadmill exercise reverses dendritic spine loss in direct and indirect striatal medium spiny neurons in the 1-methyl-4-phenyl-1,2,3,6-tetrahydropyridine (MPTP) mouse model of Parkinson’s disease.” Neurobiol Dis 63: 201–209.

Ungerstedt, U. (1968). “6-Hydroxy-dopamine induced degeneration of central monoamine neurons.” Eur J Pharmacol 5(1): 107–110.

Vastagh, C., F. Gardoni, V. Bagetta, J. Stanic, E. Zianni, C. Giampa, B. Picconi, P. Calabresi and M. Di Luca (2012). “N-methyl-D-aspartate (NMDA) receptor composition modulates dendritic spine morphology in striatal medium spiny neurons.” J Biol Chem 287(22): 18103–18114.

Villalba, R. M. and Y. Smith (2010). “Striatal spine plasticity in Parkinson’s disease.” Front Neuroanat 4: 133.

Villalba, R. M. and Y. Smith (2018). “Loss and remodeling of striatal dendritic spines in Parkinson’s disease: from homeostasis to maladaptive plasticity?” J Neural Transm (Vienna) 125(3): 431–447.

Wall, N. R., M. De La Parra, E. M. Callaway and A. C. Kreitzer (2013). “Differential innervation of direct- and indirect-pathway striatal projection neurons.” Neuron 79(2): 347–360.

Wilson, C. J. (1987). “Morphology and synaptic connections of crossed corticostriatal neurons in the rat.” J Comp Neurol 263(4): 567–580.

Zaja-Milatovic, S., D. Milatovic, A. M. Schantz, J. Zhang, K. S. Montine, A. Samii, A. Y. Deutch and T. J. Montine (2005). “Dendritic degeneration in neostriatal medium spiny neurons in Parkinson disease.” Neurology 64(3): 545–547.

Zhuang, X., J. Masson, J. A. Gingrich, S. Rayport and R. Hen (2005). “Targeted gene expression in dopamine and serotonin neurons of the mouse brain.” J Neurosci Methods 143(1): 27–32.

