## Supplementary material for "Transiently elevated dopamine triggers plasticity that preserves motor function, striatal neuron morphology, and thalamostriatal input in a mouse model of Parkinson’s disease": Table S1, Table S2, Figure S1, Figure S2, Figure S3

**Supplemental Information**

**Table S1 - KEY RESOURCES**

| REAGENT or RESOURCE | SOURCE | IDENTIFIER |
|---|---|---|
| **Antibodies** | | |
| Rabbit anti-tyrosine hydroxylase (1:1000, IHC) | Milipore | **RRID**: AB_390204 |
| AffiniPure Donkey anti-rabbit Alexa Fluor 488 (1:500, IHC) | Jackson ImmunoResearch | **RRID:** AB_2313584 |
| Streptavidin Alexa Fluor 647 (1:500, IHC) | Jackson ImmunoResearch | **RRID**: AB_2341101 |
| **Bacterial and Virus Strains** | | |
| AAV8-hSyn-DIO-hM3D(Gq)-mCherry | Addgene | 44361 |
| AAV8-hSyn-DIO-mCherry | Addgene | 50459 |
| AAV1.Syn.Chronos-GFP.WPRE | Addgene | 59170 |
| AAV5.Syn.Chronos-GFP.WPRE | Addgene | 59170 |
| AAV5/Ef1a-Flex-Chronos-GFP | UNC Vector Core | AV6474 |
| AAV1.hSyn.ReaChR-mCitrine.WPRE.SV40 | Addgene | 50954 |
| AAV1.hSyn.Flex.ReaChR-mCitrine.WPRE.SV40 | Addgene | 50955 |
| **Chemicals, Peptides, and Recombinant Proteins** | | |
| 6-hydroxydopamine hydrochloride | Sigma Life Sciences | H4381-100mg |
| Cholera Toxin Subunit B, Alexa Fluor 488 | ThermoFisher Scientific | C34775 |
| Clozapine-N-Oxide | TOCRIS Bioscience | 4936 |
| SCH23390 hydrochloride | TOCRIS Bioscience | 0925 |

| | | |
|---|---|---|
| 1,1'-Dioctadecyl-3,3,3',3'-Tetramethylindocarbocyanine Perchlorate (DiI) | ThermoFisher Scientific | D3911 |
| Neurobiotin | Vector Laboratories | **RRID**: AB_2336606 |
| AnaSed (Xylazine) | Henry Schein Animal Health | 033197 |
| Isothesia (Isoflurane) | Henry Schein Animal Health | 029405 |
| Ketoprofen (Ketofen) | Zoetis | NADA 140-269 |
| Tetrodotoxin (TTX) | Tocris | 1069 |
| CPP | Tocris | 0173 |
| 4-Aminopyridine (4-AP) | Sigma-Aldrich | 275875 |
| **Experimental Models: Organisms/Strains** | | |
| Slc6a3tm1(cre)Xz/J (DATcre) | Jackson Laboratories | **RRID:** IMSR_JAX:020080 |
| B6.Cg-Tg(Drd1a-tdTomato)6Calak/J ($D1^{dt}$) | Jackson Laboratories | **RRID:** IMSR_JAX:016204 |
| $Slc17a8^{-/-}$ ($VGLUT3^{-/-}$) | Rebecca Seal | n/a |
| Tlx3_PL56 | MMRRC | 041158-UCD |
| Sim1_KJ18 | MMRRC | 034746-UCD |
| C57Bl/6J | Jackson Laboratories | **RRID:** IMSR_JAX:000646 |
| **Software and Algorithms** | | |
| Motor Monitor | Kinder Scientific | 16224-11 |
| PRISM v8 | GraphPad | **RRID:** SCR_002798 |
| NIS-Elements | Nikon | **RRID:** SCR_014329 |
| Fiji (ImageJ) | NIH | **RRID:** SCR_002285 |
| Axon pCLAMP v10 | Molecular Devices | **RRID:** SCR_011323 |
| Matlab | Mathworks | R2020b (other versions compatible) |
| Wavesurfer | Janelia Research Campus | https://wavesurfer.janelia.org/ |

### Table S2 - Statistics

| Fig/Panel | Group | Test | N/Size | *P* value |
|---|---|---|---|---|
| 1D | Cylinder test, WT, ipsi vs contra | Mixed-effects model (REML) | N=10-13 per group | 0.0007 |
| 1D | Cylinder test, cTED, ipsi vs contra | Mixed-effects model (REML) | N=8-10 per group | 0.0271 |
| 1E | Cylinder test, WT baseline vs depleted, sleeping | Independent samples test | N=8 mice | 0.0002 |
| 1E | Cylinder test, WT baseline vs depleted, waking | Independent samples test | N=10 mice | 1.64E-9 |
| 1E | Cylinder test, cTED baseline vs depleted, sleeping | Independent samples test | N=11 mice | 0.314 |
| 1E | Cylinder test, cTED baseline vs depleted, waking | Independent samples test | N=13 mice | 0.072 |
| 1F | TH loss, WT vs. cTED | Independent samples test | N=10 per group | 0.231 |
| 2D | TH loss, WT vs. iTED | 1-way ANOVA | N=11-14 per group | 0.288 |
| 2E | 24 hr activity. mCherry vs. eDREADD sal vs. iTED | 2-way ANOVA (posthoc multiple comparisons) | N=7-10 per group | <0.0001 |
| 2F | Cylinder test, mCherry+CNO ipsi vs contra | Independent samples test | N=12 mice | p<0.0001 |
| 2F | Cylinder test, eDREADD+Saline: ipsi vs contra | Independent samples test | N=11 mice | p<0.0001 |
| 2F | Cylinder test, eDREADD+CNO: ipsi vs contra | Independent samples test | N=14 mice | 0.0719 |
| 3B | cTED Mature spine #, WT baseline vs depleted, waking | 2-way ANOVA (post-hoc Bonferroni multiple comparison) | N=3 mice and n=7 dendrites/group | p<0.0001 |
| 3B | cTED Mature spine #, WT baseline vs depleted, sleeping | 2-way ANOVA (post-hoc Bonferroni multiple comparison) | N=3 mice and n=7 dendrites/group | 0.0025 |
| 3B | cTED Mature spine #, Constitutive TED mice, baseline vs depleted, waking | 2-way ANOVA (post-hoc Bonferroni multiple comparison) | N=3-4 mice and n=9-12 dendrites/group | P<0.0001 |
| 3B | cTED Mature spine #, Constitutive TED mice, baseline vs depleted, sleeping | 2-way ANOVA (post-hoc Bonferroni multiple comparison) | N=3-4 mice and n=6 dendrites/group | 0.003 |
| 3C | D1R Mature spine #, ctl vs iTED, vs iTED-CNO baseline vs depleted, waking | 2-way ANOVA (post-hoc Bonferroni multiple comparison) | N=3-4 mice and n=6-12 dendrites/group | 0.029 |
| 3C | D1R Mature spine #, ctl vs iTED, vs iTED-CNO baseline vs depleted, sleeping | 2-way ANOVA (post-hoc Bonferroni multiple comparison) | N=3-4 mice and n=5 dendrites/group | 0.0027 |
| 3C | D2R Mature spine #, ctl vs iTED, vs iTED-CNO baseline vs depleted, waking | 2-way ANOVA (post-hoc Bonferroni multiple comparison) | N=3-4 mice and n=9-12 dendrites/group | 0.0003 |
| 3C | D2R Mature spine #, ctl vs iTED, vs iTED-CNO baseline vs depleted, sleeping | 2-way ANOVA (post-hoc Bonferroni multiple comparison) | N=3-4 mice and n=5 dendrites/group | 0.02 |
| 3D | D1R mature spine #, ctl vs cTED +antagonist, baseline vs depleted | 2-way ANOVA (post-hoc Bonferroni multiple comparison) | N=3-4 mice and n=6-9 dendrites/group | 0.04 |
| 3D | D2R mature spine #, ctl vs cTED +antagonist, baseline vs depleted | 2-way ANOVA (post-hoc Bonferroni multiple comparison) | N=3-4 mice and n=6-9 dendrites/group | P<0.0001 |

| | | | | |
|---|---|---|---|---|
| 4C | Total dendritic length mCherry, eDREADD-sal, eDREADD-cno, baseline vs depletion | 1-way ANOVA (post-hoc Tukey multiple comparison) | N=3-4 mice per grp. N=4-11 dendrites per grp. | P<0.0001 |
| 5D left | Number of Action potentials, (125mV) (WT, WT depleted, cTED, cTED depleted) | 1-way ANOVA (post-hoc Tukey multiple comparison) | WT,n=34; WT depl.,n=24; cTED,n=29; cTED depl.,n=21 neurons | 0.01(All); groups: 0.04(WT vs WT depl.); 0.04(WT vs cTED depl.) |
| 5D left | Number of Action potentials, (150mV) (WT, WT depleted, cTED, cTED depleted) | 1-way ANOVA (post-hoc Tukey multiple comparison) | WT,n=34; WT depl.,n=24; cTED,n=29; cTED depl.,n=21 neurons | 0.0002(All); groups: 0.03(WT vs WT depl.); 0.0004(WT vs cTED depl.); 0.004(cTED vs. cTED depl.) |
| 5D left | Number of Action potentials, (175mV) (WT, WT depleted, cTED, cTED depleted) | 1-way ANOVA (post-hoc Tukey multiple comparison) | WT,n=34; WT depl.,n=24; cTED,n=29; cTED depl.,n=21 neurons | <0.0001(All); groups: 0.0001(WT vs cTED depl.); 0.001(cTED vs. cTED depl.) |
| 5D left | Number of Action potentials, (200mV) (WT, WT depleted, cTED, cTED depleted) | 1-way ANOVA (post-hoc Tukey multiple comparison) | WT,n=34; WT depl.,n=24; cTED,n=29; cTED depl.,n=21 neurons | <0.0001(All); groups: 0.046(WT vs WT depl.); <0.0001(WT vs cTED depl.); 0.0004(cTED vs. cTED depl.) |
| 5D left | Number of Action potentials, (225mV) (WT, WT depleted, cTED, cTED depleted) | 1-way ANOVA (post-hoc Tukey multiple comparison) | WT,n=34; WT depl.,n=24; cTED,n=29; cTED depl.,n=21 neurons | 0.0002(All); groups: 0.0001(WT vs cTED depl.); 0.002(cTED vs. cTED depl.) |
| 5D left | Number of Action potentials, (250mV) (WT, WT depleted, cTED, cTED depleted) | 1-way ANOVA (post-hoc Tukey multiple comparison) | WT,n=34; WT depl.,n=24; cTED,n=29; cTED depl.,n=21 neurons | 0.002(All); groups: 0.001(WT vs cTED depl.); 0.01(cTED vs. cTED depl.) |
| 5D left | Number of Action potentials, (275mV) (WT, WT depleted, cTED, cTED depleted) | 1-way ANOVA (post-hoc Tukey multiple comparison) | WT,n=34; WT depl.,n=24; cTED,n=29; cTED depl.,n=21 neurons | 0.04(All); groups: 0.04(WT vs cTED depl.) |
| 5E left, (75mV) | Average firing frequency (WT, WT depleted, cTED, cTED depleted) | 1-way ANOVA (post-hoc Tukey multiple comparison) | WT,n=34; WT depl.,n=24; cTED,n=29; cTED depl.,n=21 neurons | 0.02 (All); groups: 0.02(WT depl. vs cTED); 0.04(WT depl. vs cTED depl.) |
| 5E left, (100mV) | Average firing frequency (WT, WT depleted, cTED, cTED depleted) | 1-way ANOVA (post-hoc Tukey multiple comparison) | WT,n=34; WT depl.,n=24; cTED,n=29; cTED depl.,n=21 neurons | 0.007(All); groups: 0.02(WT vs WT depl.); 0.02(WT depl. vs cTED); 0.03(WT depl. vs cTED depl.) |
| 5E left, (125mV) | Average firing frequency (WT, WT depleted, cTED, cTED depleted) | 1-way ANOVA (post-hoc Tukey multiple comparison) | WT,n=34; WT depl.,n=24; cTED,n=29; cTED depl.,n=21 neurons | 0.002(All); groups: 0.001(WT vs WT depl.); 0.01(WT depl. vs cTED); 0.04(WT depl. vs cTED depl.) |
| 5E left, (150mV) | Average firing frequency (WT, WT depleted, cTED, cTED depleted) | 1-way ANOVA (post-hoc Tukey multiple comparison) | WT,n=34; WT depl.,n=24; cTED,n=29; cTED depl.,n=21 neurons | 0.0003(All); groups: 0.0004(WT vs WT depl.); 0.001(WT depl. vs cTED) |
| 5E left, (175mV) | Average firing frequency (WT, WT depleted, cTED, cTED depleted) | 1-way ANOVA (post-hoc Tukey multiple comparison) | WT, n=34; WT depl., n=24; cTED, n=29; cTED depl., n=21 neurons | 0.0005(All); groups: 0.001(WT vs WT depl.); 0.002(WT depl. vs cTED) |
| 5E left, (200mV) | Average firing frequency (WT, WT depleted, cTED, cTED depleted) | 1-way ANOVA (post-hoc Tukey multiple comparison) | WT, n=34; WT depl., n=24; cTED, n=29; cTED depl., n=21 neurons | 0.002(All); groups: 0.002(WT vs WT depl.); 0.01(WT depl. vs cTED) |
| 5E left, (225mV) | Average firing frequency (WT, WT depleted, cTED, cTED depleted) | 1-way ANOVA (post-hoc Tukey multiple comparison) | WT, n=34; WT depl., n=24; cTED, n=29; cTED depl., n=21 neurons | 0.01(All); groups: 0.007(WT vs WT depl.); 0.048(WT depl. vs cTED) |

| | | | | |
|---|---|---|---|---|
| 5E left, (250mV) | Average firing frequency (WT, WT depleted, cTED, cTED depleted) | 1-way ANOVA (post-hoc Tukey multiple comparison) | WT, n=34; WT depl., n=24; cTED, n=29; cTED depl., n=21 neurons | 0.03(All); groups:0.02(WT vs WT depl.) |
| 5F left | dSPN: resting membrane potential | Student's 2-tailed t-test | WT, n=34; WT depl., n=24; cTED, n=29; cTED depl., n=21 neurons | 0.01(WT vs WT depl.); 0.005(WT depl. vs cTED depl.) |
| 5F right | iSPN: resting membrane potential | Student's 2-tailed t-test | WT, n=25; WT depl., n=24; cTED, n=24; cTED depl., n=21 neurons | 0.003 (WT vs WT depl.); 0.03(WT depl. Vs cTED) |
| 6C | Total EPSC amplitude from IT to dSPN and iSPN | Mann-Whitney 2-tailed test | dSPN (WT, n=28; WT depl., n=10); iSPN (WT, n=22; WT depl., n=9) neurons | 0.006 (dSPN:WT vs WT depl.); 0.03 (iSPN:WT vs WT depl.) |
| 6D | Total EPSC amplitude from IT to dSPN and iSPN | Mann-Whitney 2-tailed test | dSPN (cTED, n=25; cTED depl., n=7); iSPN (cTED, n=21; cTED depl., n=7) | <0.0001 (dSPN:cTED vs cTED depl.); <0.0001 (iSPN:cTED vs cTED depl.) |
| 6G | Total EPSC amplitude from PT to dSPN and iSPN | Mann-Whitney 2-tailed test | dSPN (WT, n=25; WT depl., n=7); iSPN (WT, n=19; WT depl., n=9) neurons | 0.007 (dSPN:WT vs WT depl.); 0.0006 (iSPN:WT vs WT depl.) |
| 6H | Total EPSC amplitude from PT to dSPN and iSPN | Mann-Whitney 2-tailed test | dSPN (cTED, n=9; cTED depl., n=4); iSPN (cTED, n=7; cTED depl., n=4) | 0.003 (dSPN:cTED vs cTED depl.); 0.006 (iSPN:cTED vs cTED depl.) |
| 7C | EPSC amplitude to dSPN, iSPN from PF input | 1-way ANOVA (post-hoc Tukey multiple comparison) | dSPN (WT,n=14; WT depl.,n=17) iSPN (WT,n=11; WT depl.,n=16) neurons | n.s. |
| 7D | EPSC amplitude to dSPN, iSPN from PF input | 1-way ANOVA (post-hoc Tukey multiple comparison) | dSPN (cTED,n=23; cTED depl.,n=15); iSPN (cTED,n=24; cTED depl.,n=15) neurons | n.s. |
| 7E | dSPN/iSPN strength ratio; EPSC amplitude from PF input | 1-way ANOVA (post-hoc Tukey multiple comparison) | WT,n=25; WT depl.,n=21; cTED,n=24; cTED depl.,n=29 neurons | 0.03 (dSPN/iSPN:WT vs cTED); 0.005 (dSPN/iSPN:WT depl. vs cTED); <0.0001 (dSPN/iSPN: cTED vs cTED depl.) |
| 7G | NMDAR amplitude to dSPN, iSPN from PF input | 1-way ANOVA (post-hoc Tukey multiple comparison) | dSPN (WT,n=12; WT depl.,n=15) iSPN (WT,n=10; WT depl.,n=13) neurons | n.s. |
| 7H | NMDAR amplitude to dSPN, iSPN from PF input | 1-way ANOVA (post-hoc Tukey multiple comparison) | dSPN (cTED,n=17; cTED depl.,n=15); iSPN (cTED,n=14; cTED depl.,n=13) neurons | n.s. |
| 7I | dSPN/iSPN strength ratio; NMDAR amplitude from PF input | 1-way ANOVA (post-hoc Tukey multiple comparison) | WT,n=20; WT depl.,n=18; cTED,n=13; cTED depl.,n=25 neurons | n.s. |
| 7J | NMDAR/AMPAR current ratio; PF input to dSPN | 1-way ANOVA (post-hoc Tukey multiple comparison) | WT,n=12; WT depl.,n=15; cTED,n=17; cTED depl.,n=15 neurons | 0.048 (WT vs WT depl.); 0.001 (WT depl. vs cTED); 0.03 (WT depl. vs cTED depl.) |
| 7K | NMDAR/AMPAR current ratio; PF input to iSPN | 1-way ANOVA (post-hoc Tukey multiple comparison) | WT,n=9; WT depl.,n=12; cTED,n=14; cTED depl.,n=11 neurons | n.s. |
| 8C | Normalized paired-pulse ratio, PR to dSPN, at 2$^{nd}$ Pulse | 1-way ANOVA (post-hoc Tukey multiple comparison) | WT,n=17; WT depl.,n=15; | 10 Hz, 0.03(All), 0.049 (WT vs WT depl.); |

| | | | | |
|---|---|---|---|---|
| | | | cTED,n=14; cTED depl.,n=14 neurons | 20 Hz; 0.01(All), 0.04 (WT vs WT depl.), 0.02(WT depl. vs cTED); 33 Hz: n.s. |
| 8C | Normalized paired-pulse ratio, PR to dSPN, at 3rd Pulse | 1-way ANOVA (post-hoc Tukey multiple comparison) | WT,n=17; WT depl.,n=15; cTED,n=14; cTED depl.,n=14 neurons | 10 Hz, 0.047(All), 0.04 (WT vs WT depl.); 20 Hz; 0.01(All), 0.04 (WT vs WT depl.), 0.01(WT depl. vs cTED); 33 Hz: n.s. |
| 8C | Normalized paired-pulse ratio, PR to dSPN, 4th Pulse | 1-way ANOVA (post-hoc Tukey multiple comparison) | WT,n=17; WT depl.,n=15; cTED,n=14; cTED depl.,n=14 neurons | 10 Hz, 0.02(All), 0.01 (WT vs WT depl.); 20 Hz: n.s.; 33 Hz: n.s. |
| 8C | Normalized paired-pulse ratio, PR to dSPN, 5th Pulse | 1-way ANOVA (post-hoc Tukey multiple comparison) | WT,n=17; WT depl.,n=15; cTED,n=14; cTED depl.,n=14 neurons | 10 Hz, 0.04(All), 0.03 (WT vs WT depl.); 20 Hz: n.s.; 33 Hz: n.s. |
| 8D | Normalized paired-pulse ratio, PR to iSPN, at 2nd Pulse | 1-way ANOVA (post-hoc Tukey multiple comparison) | WT,n=15; WT depl.,n=14; cTED,n=17; cTED depl.,n=15 neurons | 10 Hz: n.s.; 20 Hz, 0.003(All), 0.008 (WT vs WT depl.), 0.007(WT depl. vs cTED); 33 Hz: n.s. |
| 8D | Normalized paired-pulse ratio, PR to iSPN, at 3rd Pulse | 1-way ANOVA (post-hoc Tukey multiple comparison) | WT,n=15; WT depl.,n=14; cTED,n=17; cTED depl.,n=15 neurons | 10 Hz, 0.04(All), 0.04(WT depl. vs cTED); 20 Hz, 0.002(All), 0.003(WT vs WT depl.), 0.01(WT depl. vs cTED); 33 Hz: n.s. |
| 8D | Normalized paired-pulse ratio, PR to iSPN, 4th Pulse | 1-way ANOVA (post-hoc Tukey multiple comparison) | WT,n=15; WT depl.,n=14; cTED,n=17; cTED depl.,n=15 neurons | 10 Hz, 0.005(All), 0.01(WT vs WT depl.), 0.01(WT depl. vs cTED); 20 Hz, 0.001(All), 0.001(WT vs WT depl.), 0.02(WT depl. vs cTED); 33 Hz: n.s. |
| 8D | Normalized paired-pulse ratio, PR to iSPN, 5th Pulse | 1-way ANOVA (post-hoc Tukey multiple comparison) | WT,n=15; WT depl.,n=14; cTED,n=17; cTED depl.,n=15 neurons | 10 Hz, 0.01(All), 0.04(WT vs WT depl.), 0.02(WT depl. vs cTED) 20 Hz, 0.001(All), 0.002(WT vs WT depl.), 0.01(WT depl. vs cTED); 33 Hz: n.s. |

**Supplemental Figure 1 | D1R receptor activation mediates the m otor-protective effects of Transient Elevation of DA (TED)**

(A) The constitutive TED model (cTED) is used in conjunction with a D1R antagonist to test for the role of D1R in mediating the neuroprotective effects. (B) Timeline of D1R antagonist injection prior to (7 days) and following DA depletion via intrastriatal 6-OHDA injection. (C) Increased motor activity occurs during the waking cycle in saline treated cTED mice (red) but this effect is blocked by the D1R antagonist SCH23390 (purple). (D) Control mice (WT, black) and D1R antagonist treated cTED mice (purple) show significant differences in ipsilateral paw dominance post-depletion compared to cTED mice (VGLUT3 KO, red). Dotted lines indicate 50% (normal) and 70% (Parkinsonian) ipsilateral paw reaches, respectively. One-way repeated measures ANOVA with Bonferroni post hoc comparison to baseline *p<0.05, **p<0.01, ***p<0.001, ****p<0.0001, ns = not significant. (E) 6-OHDA depletions result in similar reduction in TH+ axons in striatum across conditions, as quantified by percent of striatal area lost.

**Supplemental Figure 2 | Multiple Days of Transient Elevation of Dopamine (TED) is required for preservation of motor behavior**

(A) The inducible model of Transient Elevation of Dopamine (TED) in which DAT-Cre mice express eDREADD (hM3Dq) via stereotaxic AAV injection into SNpC. (B) This permits control over the amount of time that elevated SNpC activity is needed for preservation of motor behavior. Here, a timeline showing 1, 3, or 7 days of eDREADD-mediated elevation of SNpC activity prior to DA depletion and behavioral testing. (C) Cylinder test for paw preference following unilateral 6-OHDA injection counts paw touches on control and depleted side paws, showing that 3 or 7 days but not 1 day of eDREADD-mediated elevation of SNpC activity can protect mice against loss of motor function following 1 week of unilateral DA depletion.

**Supplemental Figure 3 | Transient Elevation of DA (TED) does not preserve immature dSPN and iSPN spines following DA depletion**

(A) Immature spine data for the cTED model (VGLUT3 KO) presented as in Figure 3. Circles represent number of spines per dendrites measured. Left, long/thin spines; right, stubby spines. dSPN in top row, iSPN below. (B) Immature spine data for the inducible TED model (eDREADD+CNO) presented as in Figure 3. Left, long/thin spines; right, stubby spines. dSPN in top row, iSPN below. (C) Immature spine data for D1R antagonist experiments.

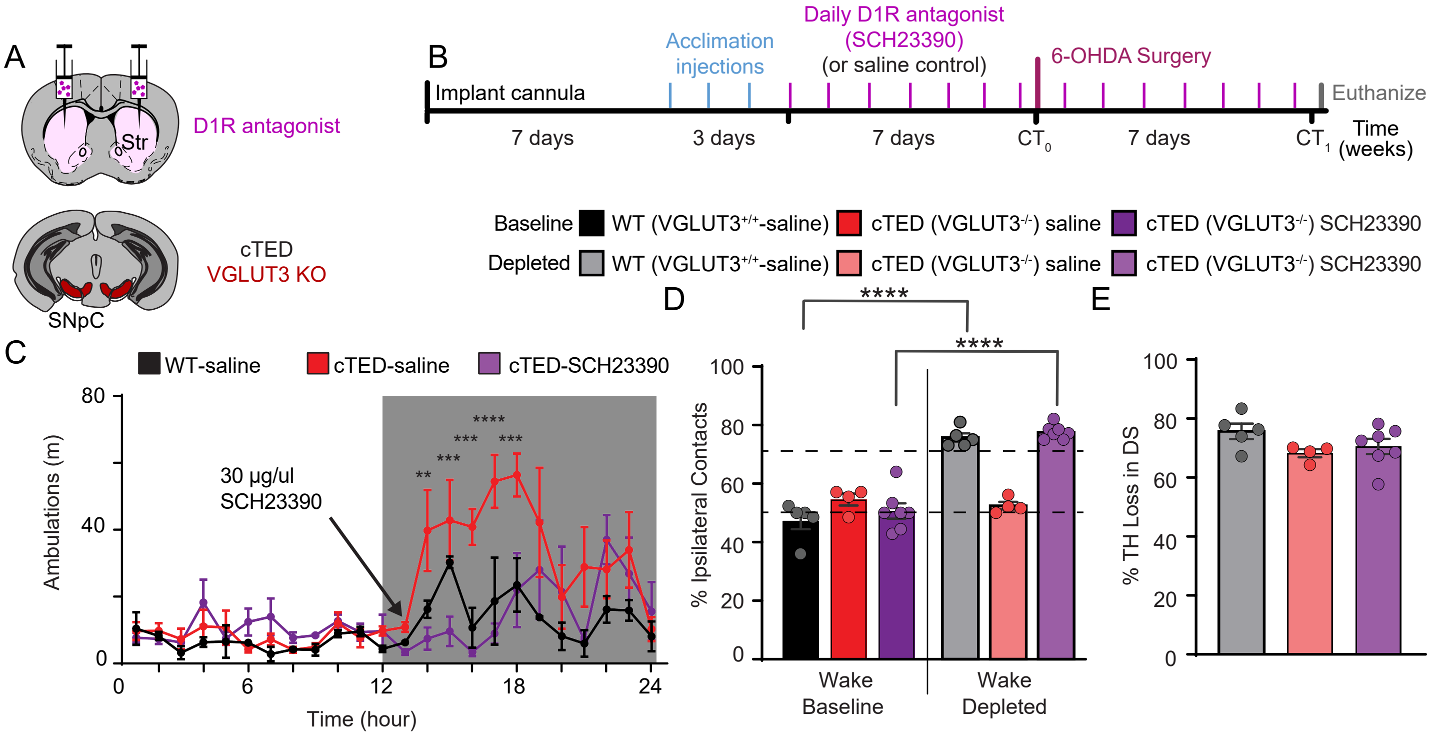

**Supplemental Figure 1 | D1R receptor activation mediates the motor-protective effects of Transient Elevation of DA (TED)**
(A) The constitutive TED model (cTED) is used in conjunction with a D1R antagonist to test for the role of D1R in mediating the neuroprotective effects. (B) Timeline of D1R antagonist injection prior to (7 days) and following DA depletion via intrastriatal 6-OHDA injection. (C) Increased motor activity occurs during the waking cycle in saline treated cTED mice (red) but this effect is blocked by the D1R antagonist SCH23390 (purple). (D) Control mice (WT, black) and D1R antagonist treated cTED mice (purple) show significant differences in ipsilateral paw dominance post-depletion compared to cTED mice (VGLUT3 KO, red). Dotted lines indicate 50% (normal) and 70% (Parkinsonian) ipsilateral paw reaches, respectively. One-way repeated measures ANOVA with Bonferroni post hoc comparison to baseline *p<0.05, **p<0.01, ***p<0.001, ****p<0.0001, ns = not significant. (E) 6-OHDA depletions result in similar reduction in TH+ axons in striatum across conditions, as quantified by percent of striatal area lost.

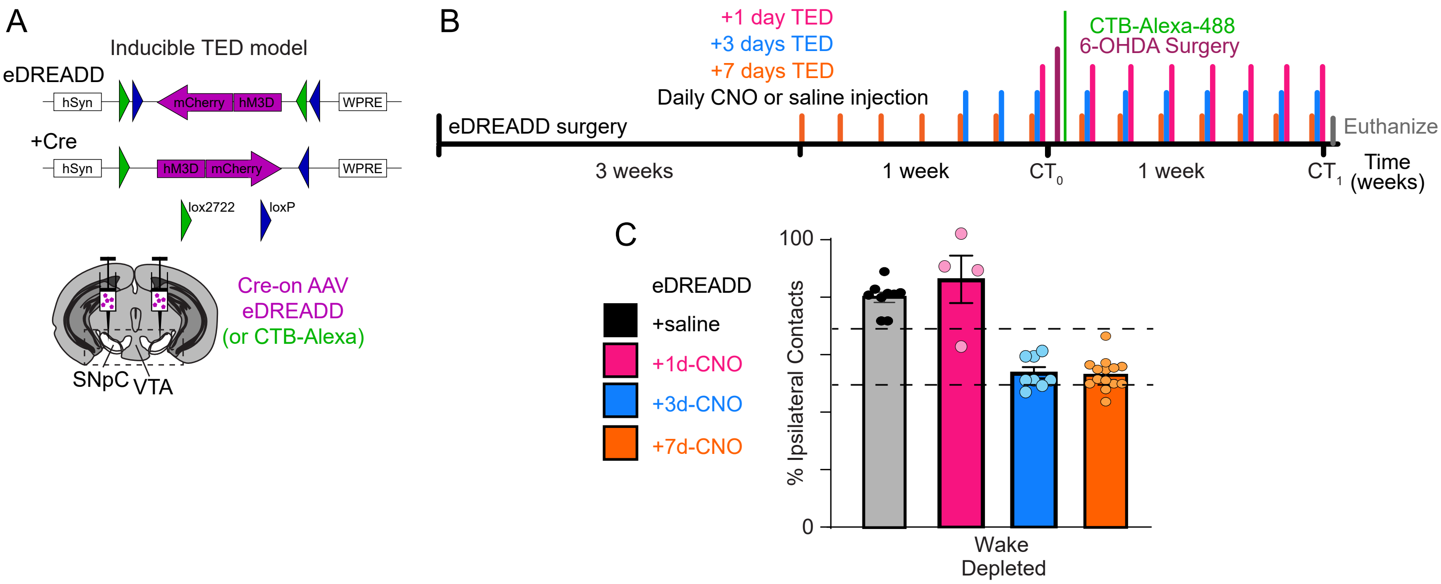

**Supplemental Figure 2 | Multiple Days of Transient Elevation of Dopamine (TED) is required for preservation of motor behavior**
(A) The inducible model of Transient Elevation of Dopamine (TED) in which DAT-Cre mice express eDREADD (hM3Dq) via stereotaxic AAV injection into SNpC. (B) This permits control over the amount of time that elevated SNpC activity is needed for preservation of motor behavior. Here, a timeline showing 1, 3, or 7 days of eDREADD-mediated elevation of SNpC activity prior to DA depletion and behavioral testing. (C) Cylinder test for paw preference following unilateral 6-OHDA injection counts paw touches on control and depleted side paws, showing that 3 or 7 days but not 1 day of eDREADD-mediated elevation of SNpC activity can protect mice against loss of motor function following 1 week of unilateral DA depletion.

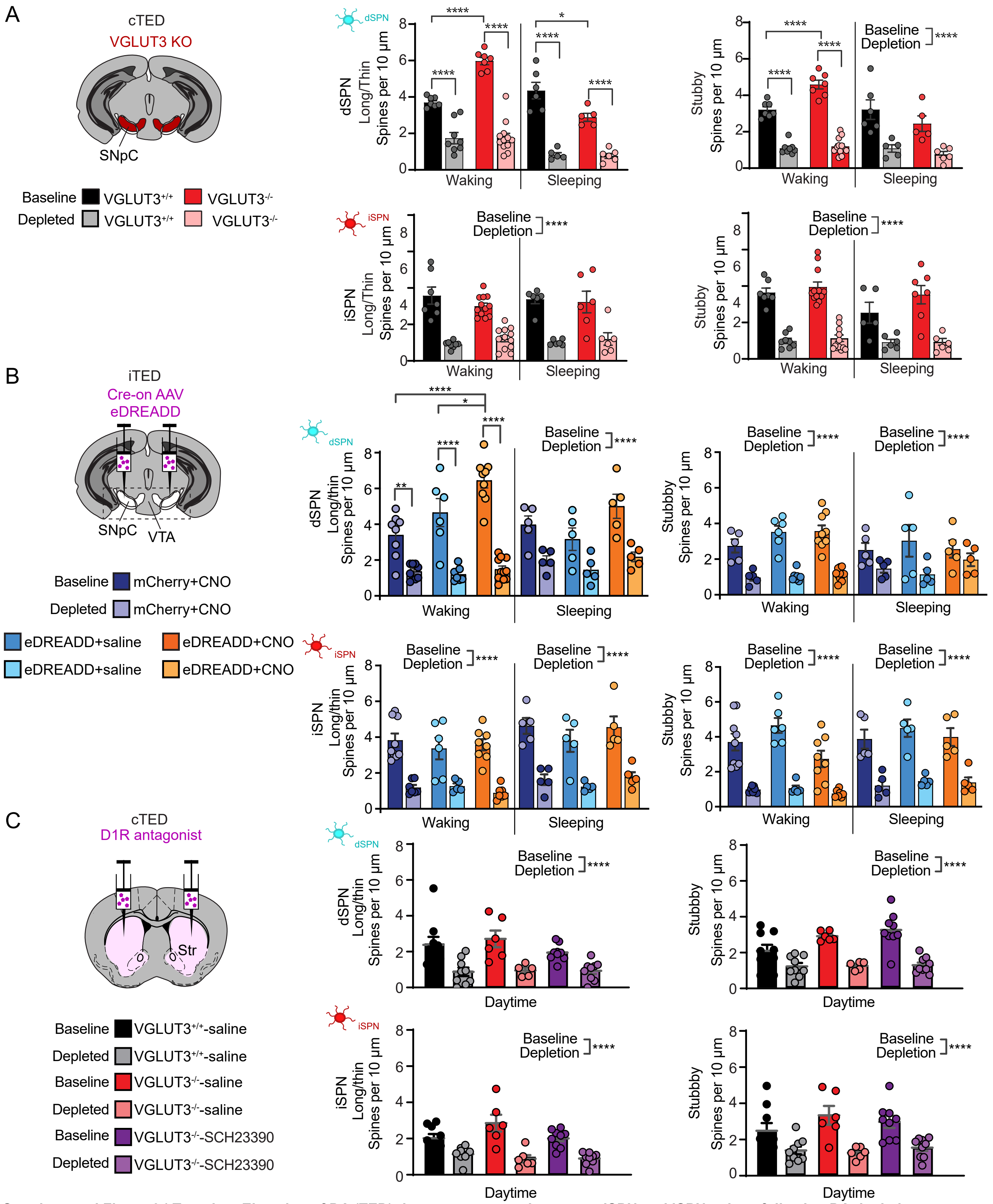

**Supplemental Figure 3 | Transient Elevation of DA (TED) does not preserve immature dSPN and iSPN spines following DA depletion**
(A) Immature spine data for the cTED model (VGLUT3 KO) presented as in Figure 3. Circles represent number of spines per dendrites measured. Left, long/thin spines; right, stubby spines. dSPN in top row, iSPN below. N=3 mice; n=7 dendrites/group. N=3 mice and n=7 dendrites/group. (B) Immature spine data for the inducible TED model (eDREADD+CNO) presented as in Figure 3. Left, long/thin spines; right, stubby spines. dSPN in top row, iSPN below. N=3-4 mice; n=6-9 dendrites/condition. N=3 mice and n=7 dendrites/group. (C) Immature spine data for D1R antagonist experiments. N=3 mice and n=6-9 dendrites/group.
